# Helical ultrastructure of the oncogenic metalloprotease meprin α in complex with a small molecule hydroxamate inhibitor

**DOI:** 10.1101/2022.03.13.484121

**Authors:** Charles Bayly-Jones, Christopher J. Lupton, Claudia Fritz, Hariprasad Venugopal, Daniel Ramsbeck, Michael Wermann, Christian Jäger, Alex de Marco, Stephan Schilling, Dagmar Schlenzig, James C. Whisstock

## Abstract

The zinc-dependent metalloprotease meprin α is predominantly expressed in the brush border membrane of proximal tubules in the kidney and enterocytes in the small intestine and colon. In normal tissue homeostasis meprin α performs key roles in inflammation, immunity, and extracellular matrix remodelling. The latter activity is furthermore important for driving aggressive metastasis in the context of certain cancers such as colorectal carcinoma. Accordingly, meprin α is the target of drug discovery programs. In contrast to meprin β, meprin α is secreted into the extracellular space, whereupon it oligomerises to form giant assemblies and is the largest extracellular protease identified to date (~6 MDa). Here, using cryo-electron microscopy, we determine the high-resolution structure of the zymogen and mature form of meprin α, as well as the structure of the active form in complex with a prototype small molecule inhibitor and human fetuin-B. Our data reveal that meprin α forms a giant, flexible, left-handed helical assembly of roughly 22 nm in diameter. We find that oligomerisation improves proteolytic and thermal stability but does not impact substrate specificity or enzymatic activity. Furthermore, structural comparison with meprin β reveal unique features of the active site of meprin α, and helical assembly more broadly.

## Introduction

Proteases critically underpin numerous processes on various levels via post-translational modification of proteins; these include cellular functions such as cell proliferation and differentiation (1, 2), necrosis (3) and apoptosis (4), angiogenesis (5), migration (6, 7) and more. Resultantly, proteases represent a fundamentally crucial component of normal cellular function and as such abnormal expression or dysregulation can be linked to various diseases. For these reasons, proteases are the attention of drug discovery programs and are exploited as diagnostic markers (8).

Astacin proteases, a subfamily of metzincin superfamily, are found in a variety of different species of vertebrates, invertebrates and bacteria. Meprin α, together with the evolutionary related meprin β, represent a subgroup of astacins found in vertebrates only (9). As with all astacins, the primary structure of the active site is characterized by two conserved motifs: the zinc-binding motif (HExxHxxGFx-HExxRxDRD) and the methionine-turn SxMHY (10, 11).

Both meprin α and β are highly expressed in epithelial cells of kidney and intestine and they have been demonstrated to a minor extent in intestinal leukocytes, skin, lung, brain, and certain cancer cells (10, 12–17). Within these tissues, numerous potential substrates are present including bioactive proteins and peptides, growth factors, adhesion molecules, and components of the extracellular matrix (ECM) (18). Differential regulation and localisation of meprins provides access to these substrates, enabling specific functions in pro- and anti-inflammatory responses, ECM assembly and remodelling, cytokine activation and signalling, as well as cell-cell adhesion (19–26).

A prominent example is the pro-collagenase function. Both meprin α and β are N- and C-pro-collagenases of procollagen I and III and are, as such, important for collagen assembly and tensile strength (19, 21, 27). Likewise, the pro-inflammatory IL1β is activated by both enzymes, although the product resulting from processing by meprin α is two-fold more active than that of meprin β (28, 29). IL6 is inactivated by meprin α and β suggesting an anti-inflammatory effect, however the soluble IL6 receptor is a product of both activities (22, 30). In contrast, pro-migratory MCP-1 and pro-inflammatory IL18 are activated by meprin α and β respectively (31, 32). These examples reflect the complexity of effects that meprins exert and participate in, with respect to pathological functions like acute kidney injury, sepsis, urinary tract infection, and inflammatory bowel disease (IBD). The roles of meprins in inflammation and modulation of immune cells was the recent topic of intensive review elsewhere (10, 25).

To achieve their function, both meprins cleave a specific but distinct motif. In meprin β this is characterised by a striking preference for negatively charged amino acids in the P1’ position unlike most other extracellular proteases (33). In contrast, meprin α favours neutral aliphatic and aromatic residues with less pronounced preference for acidic amino acids (33). As such, meprin α and β partially discriminate between substrates, resulting in distinct activity profiles. These differences are thought to drive distinct functional properties in vivo. Nevertheless, as exemplified above, some overlap in substrates also results in redundant physiological roles.

Of note, meprin α has been suggested to drive aggressive metastatic colorectal cancer (34–36). Epithelial cells of the normal human colon secret meprin α into the lumen, whereas in a colon carcinoma cell line meprin α is secreted in a nonpolarized fashion increasing substrate accessibility. Moreover, high concentrations of meprin α activity in primary tumour stroma promotes tumour progression and metastasis. This is thought to occur due to meprin α pro-migratory and pro-angiogenic activity by way of ECM remodelling and transactivation of the EGFR/MAPK signalling pathway by shedding the ligands EGF and TGFα (37). Interestingly, inhibition of meprin α by actinonin, a naturally occurring inhibitor, was able to abrogate these effects *in vitro*, suggesting meprin α is a promising target for therapeutics (37). Importantly meprin α dysregulation does not necessarily impact meprin β, whose important regulatory and physiological functions are maintained. For example, meprin β shows a protective effect in IBD and promotes mucus turnover by cleavage of MUC2, preventing bacterial overgrowth in intestinal mucosa (31, 38, 39).

Thus, non-specific inhibition of meprins may have either beneficial or deleterious effects on health depending on the context. For these reasons, specific inhibitors of meprin α have been sought to mitigate its function in disease progression and to investigate the pathophysiological role of meprins in further detail. In general, specific inhibitors for meprin α and meprin β separately might be useful to treat certain diseases, such as progressive cancers, without disrupting the physiological function of the corresponding homologue. In order to provide controlled inhibition of meprin α functions while maintaining the physiological function of meprin β, and vice versa, significant effort has been directed to develop protease specific compounds (40–42). However, owing to high conservation between the two enzymes this has been non-trivial and thus off-target inhibition of proteolytic activity is a current limitation of existing candidates.

Both meprin α and β are expressed as glycosylated zymogens. The domain architecture of both proteases comprises an N-terminal protease domain (M12A), a MAM (meprin, A-5 protein, and receptor protein-tyrosine phosphatase μ) domain, a MATH (Meprin and TNF receptor-associated factor [TRAF] homology) domain, an EGF-like domain, and, lastly, a short transmembrane and cytosolic region (Supp Figure S1a). Both proteases are enzymatically processed to remove the autoinhibitory N-terminal prodomain by trypsin-like activities (for example by the fibrinolytic protease, plasmin) in order to form the active protease (43).

A key difference between meprin α and meprin β is that the former includes a furin cleavage site between the MATH and EGF-like domain. This permits furin-mediated shedding of meprin α within the secretory pathway and, subsequently, secretion into the extracellular milieu (44, 45). In contrast, meprin β remains predominantly membrane associated, with rare shedding events mediated via ADAM10/17 proteolytic cleavage of the sequence immediately preceding the transmembrane domain (17, 18, 46, 47). Shedding of meprin β has been observed to drive distinct substrate specificity and function suggesting localisation affects accessibility to certain substrates (48). Furthermore, expression of meprin α is localised in the stratum basale of the epidermis, while meprin β is found in the stratum granulosum (15). As such, some functions of meprin α and meprin β may be dependent on their localisation and tissue distribution.

In regard to quaternary structure, both enzymes form covalently (disulphide) linked homo- and heterodimers (10). The quaternary complex is critical for the localisation and oligomerization of meprin α. Heterodimeric meprin α/β remains tethered at the plasma membrane via membrane bound meprin β (44). Such heterodimers can further interact to form heterotetramers (48–50). In contrast, furin-mediated release of meprin α homodimers results in the formation of meprin α oligomeric assemblies, as has been shown for recombinant rat meprin α (50). The non-covalent interface that mediates this homo-oligomerisation was mapped by cross-linking mass spectrometry to the MAM domain (51). In contrast, meprin β homodimers do not oligomerise when tethered to or upon shedding from the membrane (10).

In addition to low stoichiometric assemblies, the homodimer of meprin α is capable of forming large non-covalently associated soluble oligomers with a reported size of up to 6 MDa (50). Meprin α is, therefore, the largest known secreted protease complex. The oligomers have been characterized by means of SEC, EM and light scattering to be comprised of a heterogeneous population of ring, circle, spiral, and tube-like structures (50). While the crystal structure of meprin β has been solved in the active, inactive and inhibitor bound states (53, 54), the structure of meprin α remains to be determined.

The heterogeneity of meprin α oligomers precludes structure determination by X-ray crystallography, accordingly, we used cryo-EM to determine its structure. Our data show that both the zymogen and active form of the meprin α ectodomain (i.e., the region released through furin cleavage) is capable of self-association to form a flexible, giant helical assembly. Using a single particle like approach we determined the high-resolution reconstructions of meprin α in its zymogen and active state. Additionally, we determined structures of meprin α in the presence of a prototype selective inhibitor and the native inhibitor human fetuin-B. This structure represents a valuable tool for rational drug design and provides a basis for structural comparisons between the homologous enzymes. Lastly, we propose possible mechanisms for further investigation that may implicate meprin α polymerisation as an important component in correct function and regulation.

## Results

### Meprin α forms a giant flexible left-handed helical filament

It has previously been observed that meprin α function requires two proteolytic events. Firstly, the zymogen is proteolytically shed from the membrane via furin proteases, thus releasing these into the extracellular milieu. After shedding from the membrane, meprin α dimers associate non-covalently to form oligomers in the MDa range. Secondly, the zymogen is converted to the active state by proteolytic removal of the pro-peptide. To characterise the structural and biochemical properties of meprin α, we expressed a truncated recombinant construct in S2 cells that lacked both the EGF-like and transmembrane regions (Supp Figure S1a, b). The secreted wild type meprin α was purified from the conditioned media via hydrophobic interaction followed by affinity chromatography (Supp Figure S1c, d). Since size exclusion chromatography regularly resulted in loss of protein and activity this method was avoided. Activation of the zymogen was accomplished by way of magnetic trypsin beads. These enabled the cleavage of the pro-peptide and facilitated subsequent removal of trypsin without further dilution of the protein sample by additional purification steps.

Inspection of the recombinant material by electron microscopy revealed meprin α forms extraordinarily large and flexible helical filaments (Figure 1a, Supp Movie 1, Supp Table 1). To obtain the structure of these filaments, we resorted to a single particle-like analysis to overcome the intrinsic flexibility and symmetry breaking *in silico* (Supp Figure S2). In this approach, ~20 nm segments of the filament were reconstructed without symmetry generating a 10–12 Å resolution reconstruction, which suffered from errors in alignment due to global flexibility. This reconstruction enabled nonoverlapping sub-regions consisting of two meprin α dimers to be localised within the original extracted particles with roughly nanometre precision. High resolution refinement of these sub-regions (to 2.4–3.4 Å resolution) was achieved by treating these as independent single particles after masking and subtracting the signal of neighbouring segments, thereby overcoming the prohibitive alignment errors resulting from continuous conformational heterogeneity of the filament.

**Fig. 1.**
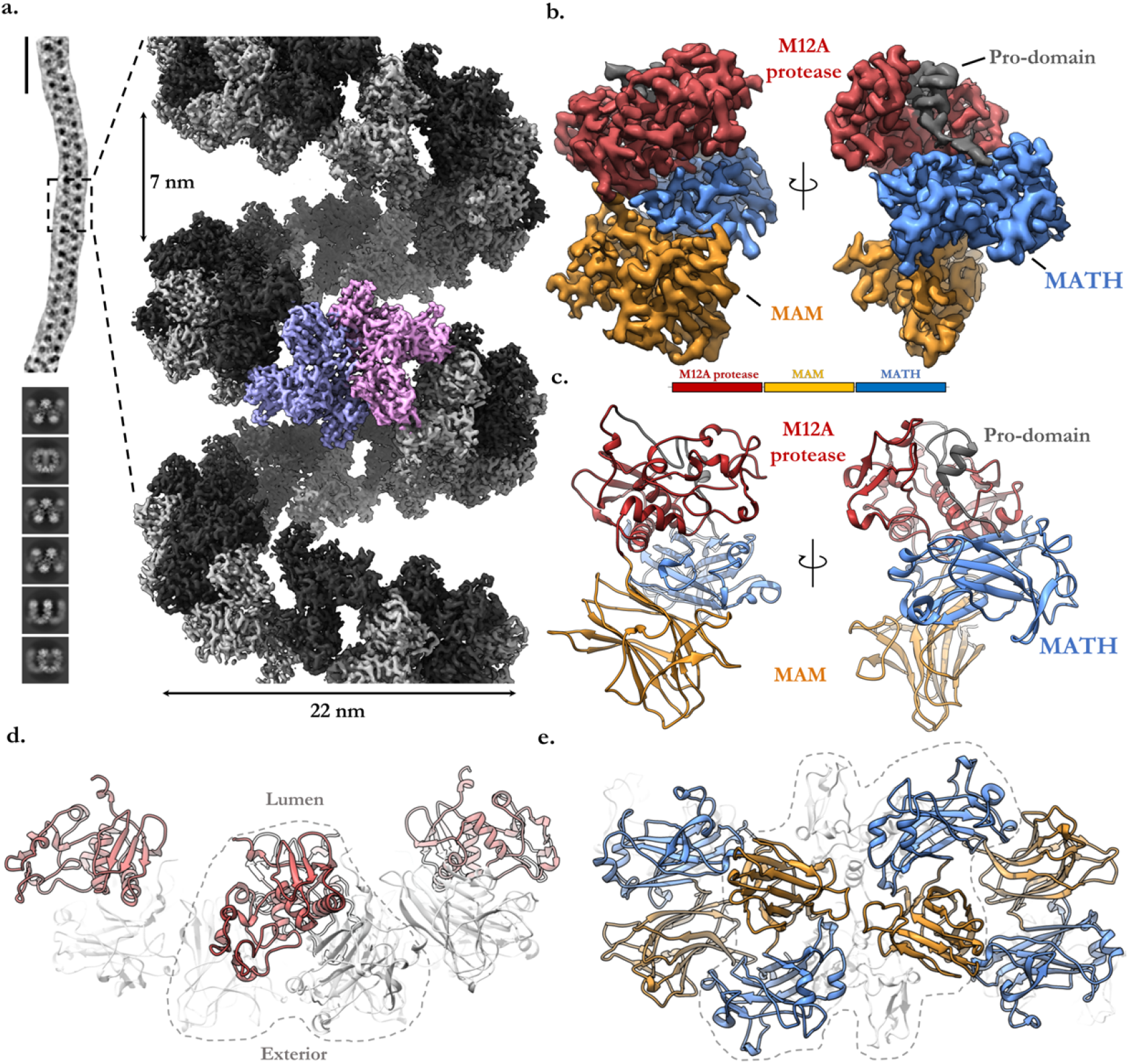
The cryo-EM reconstruction of helical meprin α. **a.** A representative particle of meprin α is roughly 100–150 nm in length (vertical scale bar, ~50 nm) and corresponding two-dimensional class averages of segmented meprin α particles. Dashed box shows the magnified view of the reconstructed segment. Homodimeric meprin α subunits are coloured alternating in black and grey. A central dimeric subunit is shown in purple and pink. **b.** A single meprin α subunit illustrating the three globular domains (M12A [red], MAM [blue] and MATH [orange]) and the helical pro-domain (dark grey). **c.** Corresponding atomic model of a single meprin α subunit and domain schematic coloured as in (b). **d.** View along the filament axis of the meprin α helix showing a segment of four meprin α subunits. The catalytic M12A domain (red) is found on the inside of the helical tube (lumen), while the MAM and MATH domains (both light grey) are located on the periphery of the helix. **e.** The meprin α helix interface is defined by two head-to-tail dimers of the MAM and MATH domains which join adjacent meprin α homodimers. In both (d) and (e) a dashed line outlines a single meprin α covalent homodimer.

The meprin α oligomer resembles a spring of roughly 22 by 200 nm in dimensions (Figure 1a, Supp Figure S3), with some filaments observed to be greater than 500 nm long (~27 MDa). We note that filaments of actomyosin (55) and the microtubule (56) possess extensive contacts between neighbouring turns. In contrast, the meprin α filament lacks such contacts, forming a highly spacious helical groove of roughly ~7 nm and a hollow core of ~15 nm (Figure 1a). Resultantly, meprin α is intrinsically flexible, observed to expand, contract, and bend along the axis of the helix (Supp Figure S3a; Supp Movie 2).

Each meprin α subunit strongly resembles the homologous protein meprin β. The subunit is a compact triad comprising the globular M12A protease, MAM and MATH domains (Figure 1b, c). In a single asymmetric unit of the helix, two meprin α subunits homodimerise into a C2 arrangement via the M12A and MATH domains. The meprin α M12A domain is typical, containing the catalytic zinc ion within the active site. Each M12A domain is positioned on the inner surface of the helix, jutting inwards forming a triangular arrangement (Figure 1d). The MAM and MATH domains buttress the protease domain, forming the outer structure of the helix interacting via a head-to-tail arrangement (Figure 1e), while at the dimer interface a conserved disulphide bond stabilises MAM/MAM interactions. This arrangement of dimers forms an indefinite filament which coils into a lefthanded helical ultrastructure.

Cisplatin treatment is known to cause an increase of meprin α in urine by inducing damage to epithelial cells in the kidney (57). To determine whether higher-order oligomers could be found within native source material, we conducted an analytical size-exclusion chromatography analysis of mouse urine after cisplatin treatment. Indeed, we observed evidence that large meprin α oligomers do exist in the urine of mice treated with cisplatin, albeit these were smaller than recombinant wild type meprin α (Supp Figure S1e).

### Analysis of meprin α oligomerisation

The assembly of meprin α is apparently governed by two unique interfaces (Figure 2a, b). The dimer interface is most extensive (1374 Å^2^) and highly conserved between both meprin α and meprin β (Figure 2c, Supp Figure S4, S5). This interface is defined by the M12A protease and MAM domains, which are dependent on a conserved MAM/MAM disulphide bond to form the covalently linked homodimer (Figure 2d). Two identical MAM/MATH interfaces (610 Å^2^) each provided by two homodimers together drive the head-to-tail formation of the non-covalent helical assembly of multiple homodimers (Figure 2a, b). This latter interface is defined by extensive charge complementarity and salt bridges between the MAM and MATH domains and has a calculated cumulative solvation free energy (Δ_i_G) of −5.2 kcal mol^−1^.

**Fig. 2.**
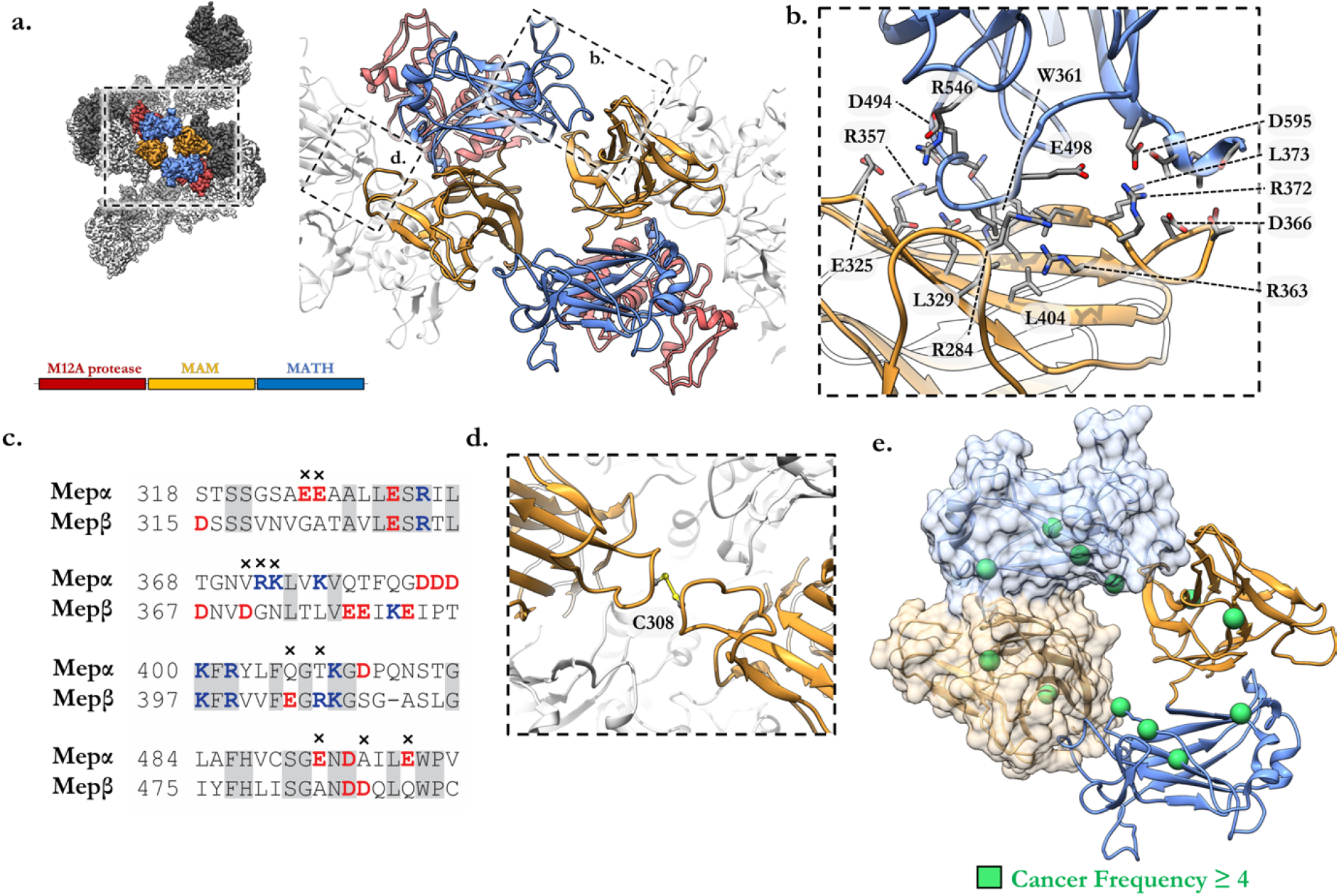
Meprin α helices are defined by two key interfaces. **a.** Segment of meprin α helix showing a single helical interface (blue, orange) defined by the MAM and MATH domains. High magnification view of this region is shown (right) as a cartoon model. **b.** Boxed region of the MAM/MATH interface illustrating key residue contacts forming at the helical interface. Visible is R372 which forms key salt bridges with adjacent aspartic acids. **c.** Local sequence alignment of MAM/MATH domains between meprin α and meprin β showing multiple charge swap and charge loss mutations. **d.** Meprin α homodimer interface showing covalent C308 disulphide bridge. **e.** Frequently occurring mutation associated with cancers (curated from the COSMIC database (52) are shown as green spheres. These residues map to the MAM/MATH domains which define the helical interface.

In contrast to meprin α, shed meprin β does not form higher-order oligomeric species. Superposition of MAM/MATH domains of meprin β and meprin α reveal the helical interface is abolished in meprin β due to multiple charge swap and loss-of-charge mutations (Figure 2c). Furthermore, multiple non-conserved residues within the MAM groove result in steric effects that likely further hinder formation of a stable interface (Figure 2c). In this regard, we note that frequently occurring cancer-associated mutations cluster to the MAM/MATH domains and, therefore, the interface of meprin α helices (Figure 2e, Extended Data 1) (52). In contrast, meprin β cancer-associated mutations more frequently cluster within the M12A protease domain (Extended Data 1) (52).

To assess the impact of these interfaces, we produced three mutants of meprin α by abolishing a salt bridge found at the helix interface (R372T and R372A) and the intermolecular disulphide bridge that stabilises the homodimer interface (C308A). No discernible differences could be observed between wild-type, R372T, and R372A meprin α by reducing and non-reducing SDS-PAGE, whereas mutant C308A appeared to be monomeric in all conditions (Supp Figure S1c, d). MALDI-TOF mass spectrometry (MS) determined the molecular mass of meprin α C308A to be exclusively monomeric (single charged ion peak at 75 kDa), while an additional disulphide bridged dimeric species (~ 130 kDA) was observed for both wild-type and R372T meprin α (Supp Figure S6a). For all variants, the zymogen form was roughly 6 kDa higher in mass than the activated counterpart consistent with the removal of the propeptide. Further, all forms had a molecular mass that exceeded the calculated mass by ~8 kDa, most likely corresponding to extensive glycosylations as observed in the cryo-EM reconstruction.

Further analysis by multi angle dynamic light scattering (MADLS) revealed different hydrodynamic radii for meprin α C308A and R372T variants, indicating the covalent and non-covalent interfaces were disrupted by mutagenesis (Supp Figure S6b). Wild type meprin α itself is polydisperse (hydrodynamic radius of main fraction ~69.6 nm), suggesting the presence of large oligomers consistent with the cryo-EM reconstruction. The observed hydrodynamic radii of C308A and R372T (~34.9 nm and ~20.8 nm respectively; Supp Figure S6b) are consistent with small oligomers or dimers being formed via the remaining unaffected interfaces. SEC-MALS analysis of C308A and R372T gave similar predictions of molecular mass of 150 kDa and 150-175 kDa respectively (Supp Figure S6c). Unlike C308A, meprin α R372T also showed some evidence of higher order oligomers under MADLS and SEC-MALS, potentially due to weak non-covalent interactions at the helix interface (Supp Figure S6c). R372A was not analysed by SEC-MALS or mass spectrometry.

Finally, inspection of variants C308A, R372T, R372A, and the dimeric meprin β homolog by electron microscopy confirmed the disruption of key interfaces. These data revealed the occurrence of small particles of dimeric appearance for R372A, while R372T and C308A were capable of homo-oligomerising into larger species, both distinct from those of wild type meprin α (Supp Figure S6d). Taken together, meprin α R327T appears to be a covalently linked dimer that has some low-affinity tendency to interact via the helical interface while R327A completely abolishes the helix interface. Conversely, C308A destabilises the major homodimer interface and forms elongated species via the helical interface and weakly via the dimer interface, consistent with previous observations (51).

### Activation and substrate specificity of meprin α

Inspection of the ultrastructure reveals the active site of the protease domain is positioned symmetrically on each edge of the helical filament, with repeating sites spaced by roughly 8 nm that line the groove of the helix (Figure 3a). Collectively, the ordered arrangement of available active sites defines a massive platform that suggests some functional purpose. We therefore sought to understand the importance of the helical assembly on meprin α substrate specificity, activity, and stability by comparing wild type meprin α to the variants, C308A, R327T and R372A.

**Fig. 3.**
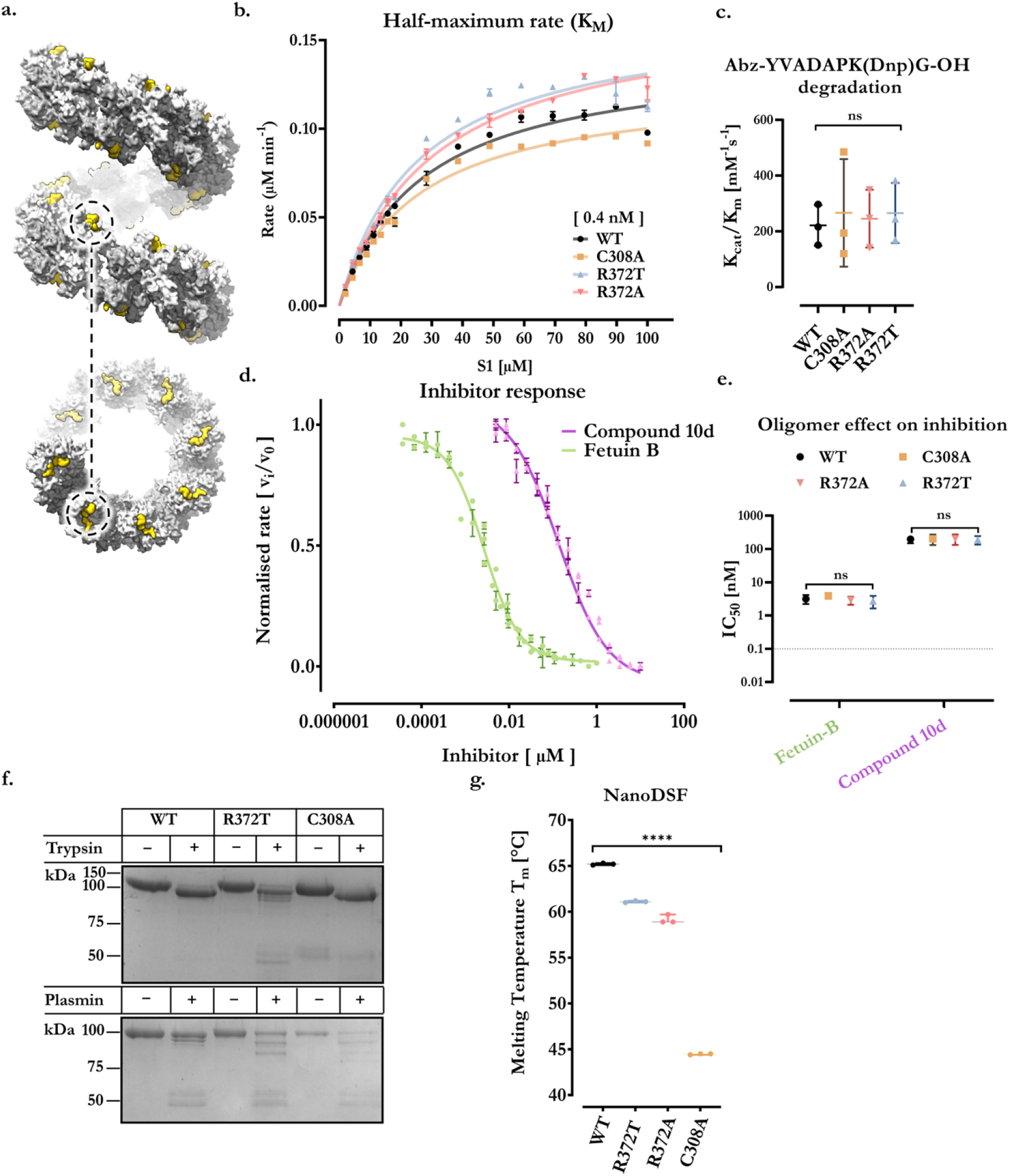
Meprin α oligomers are proteolytically and thermally stable compared to lower-stoichiometry variants. **a.** Periodic arrangement of the active site and prodomain are highlighted as yellow in the surface representation of an idealised meprin α helical segment. **b.** Example curve representing a series of first-order rates (determined by fitting the linear region of fluorescence versus time) at a given meprin concentration. The kcat/Km is determined from this graph for several meprin concentrations and in triplicate **c.** First order rate constants (k_cat_/K_m_) of meprin α and variants for small fluorogenic peptide cleavage. Oligomeric state does not appear to affect rate of cleavage of small molecule substrate. **d.** Inhibitory constant (IC_50_) of meprin α inhibitors. Determined by fitting and normalising the linear region of fluorescence versus time, in the presence of varying amount of inhibitor at a fixed meprin α concentration. **e.** Globular proteinaceous inhibitor, murine fetuin-B, and small molecule compound 10d were unaffected by oligomeric state. **f.** Proteolytic stability of meprin α and variants against trypsin and plasmin. Meprin α oligomers were more stable compared to lower-stoichiometric variants. **g.** Meprin α thermal stability measured by nanoDSF. Meprin α possess superior thermal stability compared to variants that lack either the disulphide bridge (most drastic) or helical interface interactions. All measurements are statistically significant to all others (p<0.0001, ****) unless otherwise indicated.

Thus far, comparisons between the oligomeric and dimeric meprin α *in vitro* reveal no significant difference in specific activity nor substrate specificity. When comparing the degradation turnover of large substrates by meprin α and variant C308A no dramatic differences could be observed, except for tropoelastin degradation which appears somewhat enhanced by C308A (Supp Figure S7). Similarly, the parameters for substrate hydrolysis of a small fluorogenic peptide substrate are not impacted (Figure 3b, c). Likewise, inhibition of meprin α by small molecule inhibitors are not impacted by the oligomeric state (Figure 3d, e). These findings suggest allosteric effects of the oligomer do not influence function of the catalytic core. In contrast however, dimeric meprin α was found to be less stable than the oligomeric form against proteolytic degradation by trypsin and plasmin during activation of the zymogens (Figure 3f). Furthermore, the non-oligomerising variants of meprin α showed a lower thermal stability when compared to oligomeric wild type meprin α (Figure 3g). Therefore, oligomerisation may play a role in protein regulation by way of increasing half-life in vivo.

Unlike small molecule substates and inhibitors, which are not sterically occluded by the helical packing, we investigated the inhibition of meprin α in vitro by murine fetuin-B, a 42 kDa globular protein. Remarkably, we observed no significant difference in the inhibitory capacity of fetuin-B between oligomeric and monomeric variants of meprin α indicating the oligomer does not interfere with binding (Figure 3d, e). Molecular docking analysis of fetuin-B to the meprin α oligomer instead suggests a compact, intercalated packing is possible that is not impacted by the helical rise of meprin α (Supp Figure S8a-d). Further, the interfaces predicted to form this meprin α/fetuin-B oligomer are conserved (58, 59) (Supp Figure S8f). This intercalated packing of fetuin-B is suggestive of possible higher order inhibitory oligomers. We therefore imaged meprin α in complex with human fetuin-B by cryo-EM and generated a 3.7 Å resolution reconstruction (Supp Figure S8g, Supp Figure S9). Indeed, this structure confirms the intricate packing suggested by the molecular docking analysis, albeit with some degree of flexibility. The presence of a conserved interface suggests the oligomeric nature of meprin α confers a selection pressure on fetuin-B and supports the notion that meprin α oligomers exist in vivo.

### Inhibition of meprin α

Comparison between the cryo-EM reconstructions of the zymogen and activated states reveal subtle conformational changes about the M12A catalytic domain (Figure 4a-c, Supp Table 1, Supp Figure S9). The propeptide forms a small helical cork which plugs the active core and thus sterically occludes the active site (Figure 4d). Cleavage of an exposed flexible loop, defined by residues L61-L67 located distal to the active site, releases the pro-peptide and exposes the catalytic core of the active site. Accordingly, the steric occlusion of the active site represents the major autoinhibitory mechanism. Upon activation, the two lobes of the M12A catalytic domain collapse around the active site (RMSD 3.3 Å), permitting the formation of a more tightly folded active site core (Figure 4e).

**Fig. 4.**
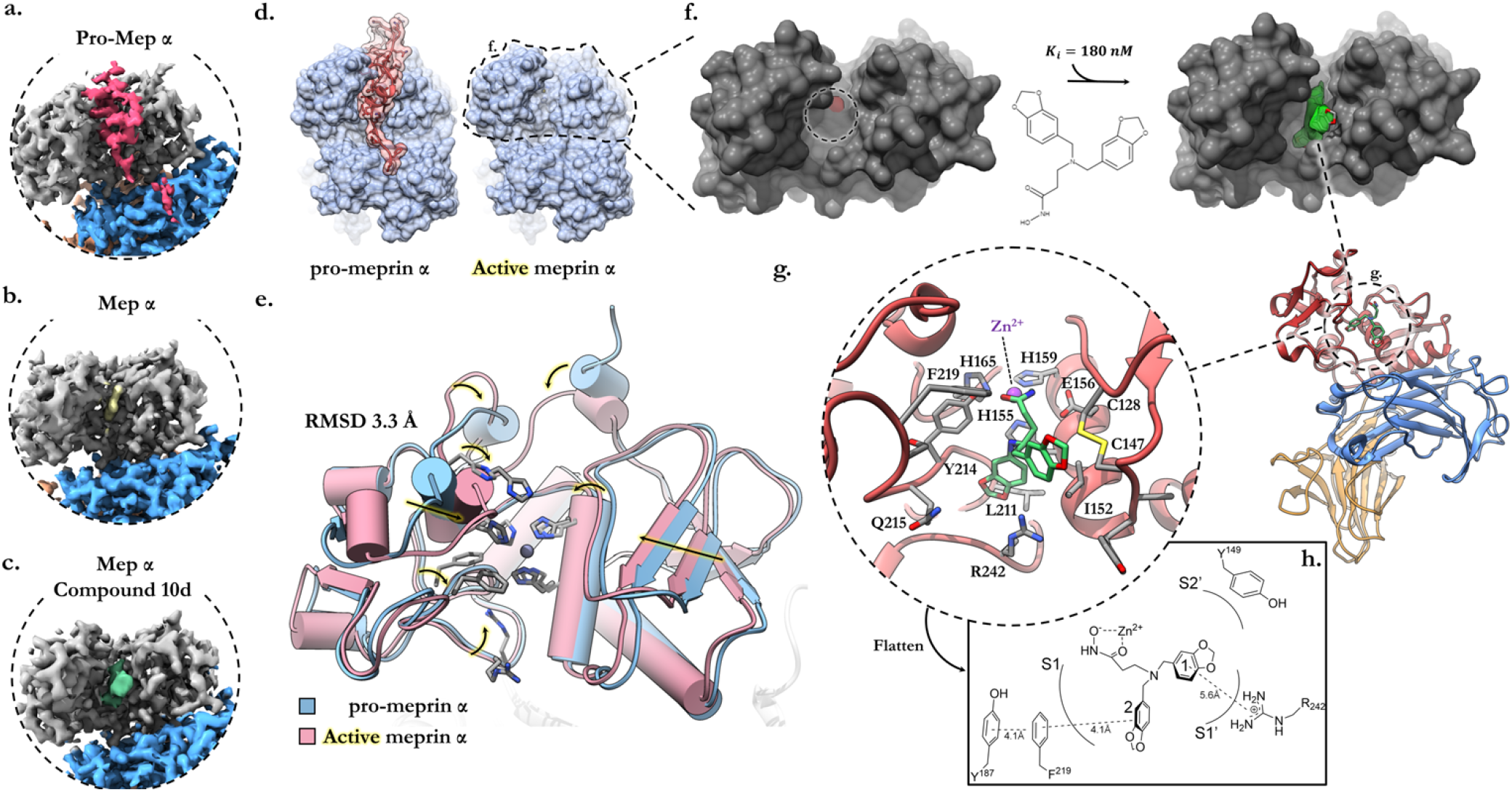
Structural basis of meprin α auto-inhibition, small molecule inhibition and activation. Zoomed view of the cryo-EM reconstructions showing **a.** the zymogen (pro-domain in red), **b.** activated form (active site in yellow) and **c.** drug inhibited form (compound 10d prototype drug in green). The M12A protease domain is coloured light grey, with the MATH domain in blue. **d.** Surface representation of the pro and active form of meprin α focused on the active site cleft. **e.** Meprin α activation requires cleavage of the pro-domain helical plug. Activation of meprin α is accompanied by conformational relaxation of the M12A protease domain resulting in a tightly folded active site core. **f.** Surface representation of meprin α M12A protease domain highlighting (dashed circle) the active site and depth of active groove. The cryo-EM density of compound 10d is shown in green, which is deeply buried within the cleft. **g.** Focused view of the meprin α active site with the small molecule inhibitor docked in its inhibitory conformation. Interactions between the hydroxamate group and zinc core constitute the major steric mechanism for inhibition of the catalytic glutamic acid (E156). The 1,3-benzodioxole groups are positioned within the deep groove forming transient interactions with aliphatic, hydrophobic and backbone functional groups. **h.** Projected two-dimensional plot of the active site and compound 10d interactions.

The majority of known meprin inhibitors possess a hydroxamic acid moiety that targets the central zinc atom of the active site (60). Thus previous generations of inhibitors, e.g. actinonin or sulphonamid-based compounds, exhibit poor specificity and can lead to off-target effects by inhibiting other metalloproteases. In particular, there is a need for specific inhibitors of meprin proteases that target either meprin α or meprin β. Our group has previously developed a small molecule inhibitor of meprin α, compound 10d (3-[bis(1,3-benzodioxol-5-ylmethyl)amino]propanehydroxamic acid) (42). The compound also binds to the zinc via a hydroamate moiety, but is built on a modified scaffhold, i.e. a tertiary amine functionalised with two 1,3-benzodioxole groups. This modified scaffhold leads to a high selectivity over other zinc-metalloproteases, i.e. MPPS and ADAMs. Compound 10d shows promising inhibitory activity and furthermore is specific to meprin α with an IC50 of 160 nM versus 2950 nM for meprin β.

To characterise the structural basis of binding and mechanism of inhibition by the small molecule inhibitor we determined the structure of meprin α after co-incubation with compound 10d. Inspection of the active site revealed new cryo-EM density positioned deeply within the groove of the active site (Figure 4c, f). Chemical docking simulations were used to flexibly fit the ligand structure to the density, revealing interactions between the hydroxamic acid group and putative zinc cation core of the catalytic triad as expected (Figure 4f, g). The two 1,3-benzodioxole groups provide further supporting interactions, predominantly from hydrophobic interactions. One benzodioxole group is deeply buried within the S1’ pocket forming an anchor point, while the other sits flush against the central helix of the M12A active site. Notably, due to the large width of the M12A groove, which accommodates the entire pro-domain α-helix, relatively few contacts are present between the M12A lobes and compound 10d. As such, we observe some degree of inhibitor mobility as assessed by loss of ordered cryo-EM density. Nevertheless, in this position both the catalytic zinc and binding pocket are sterically occluded by compound 10d, thereby inhibiting the proteolytic function of meprin α (Figure 4g).

Despite the selectivity of compound 10d for meprin α most contacts of the interaction are strongly conserved between meprin α and meprin β (Supp Figure S10). Notably, however, R238 of meprin β (R242 in meprin α) adopts a distinct rotamer position relative to meprin α, due to charge repulsion with R146 (Y149 in meprin α). In this position, R238 of meprin β sterically occludes the binding of compound 10d (Supp Figure S10). Further modification of compound 10d to exploit this charged residue and to increase contact surface area with the M12A catalytic groove may be beneficial in terms of improving affinity and specificity to meprin α.

## Discussion

Proteases are crucial players in fundamental cellular processes in human health such as proliferation, differentiation, inflammation, and ECM homeostasis, which drive various pathophysiological states when dysregulated. For example, degradation of the ECM is a typical hallmark of aggressive metastatic cancers by enabling cell migration. Meprin proteases function in the epithelium to regulate ECM homeostasis while having both anti- and pro-inflammatory roles in cell signalling (26). Indeed, meprin α has been implicated as an oncogenic factor which, when highly expressed, is correlated with more metastatic and aggressive forms of certain colorectal cancers. Further elevated levels of meprin α are observed in connection with nephritis (61–63).

Current available drugs to meprin α bind with high affinity but lack high specificity. They also inhibit the functions of meprin β. Therefore, efforts to engineer more selective and potent meprin α inhibitors are ongoing. While the structure of meprin β has been described previously, the structure of meprin α was unavailable. Structures of meprin β have provided mechanistic insights and supported rational design efforts to develop specific inhibitors to this homologue. Conversely, since the structural differences of these proteases underpin the selectivity of compounds, the absence of a meprin α structure has obfuscated drug discovery programs.

Previous studies report meprin α forms large, heterogeneous oligomeric species. We therefore used an electron microscopy approach to elucidate the structure. The high-resolution structures of the meprin α oligomer were determined in four key states. The zymogen form of meprin α is plugged by an α-helical pro-peptide that sterically occludes the active site. Upon proteolytic cleavage of the propeptide the active is exposed and the M12A protease domain undergoes a conformational relaxation. The exposed active site defines a deep pocket that positions the catalytic glutamic acid in close proximity to the substrate peptide as described for meprin β (53). Structural studies of meprin α in complex with a prototype selective inhibitor revealed the mode of inhibition. Furthermore, these data reveal subtle differences between meprin α and meprin β that may underpin inhibitor specificity. As expected, the prototype inhibitor was observed to competitively bind to the active site thereby blocking substrate engagement. These data provide long sought-after details of the meprin α active site for drug discovery programs and offer insight into the unusually large oligomeric form of meprin α.

Intriguingly, meprin α dimers associate to form giant left-handed helical filaments of seemingly no restriction to length *in vitro*. Protein oligomerisation drives many biological processes such as allosteric control of activity, regulation of protein activity by spatial sequestration, control of local concentrations, increased stability against denaturation (64, 65), and more. As such, formation of oligomers offers functional, genetic, and physicochemical advantages over monomeric counterparts. A prominent example of this includes oligomeric tripeptidyl peptidase II (TPPII), a cytosolic dimeric enzyme classified as the largest peptidase complex so far (66). But, while TPPII forms a spindle-like rigid structure, the meprin α helix is flexible with no further contacts between neighbouring turns. In TPPII this rigid structure results in restricted access to the active site and the peptidase acts only on smaller und unfolded substrates and was shown to have 10-fold increased activity after oligomerization. Overall, *in vitro* no such correlation could be found for oligomeric meprin α compared with lower-stoichiometric variants indicating that the functional role of the oligomerisation, if any, may only be apparent *in vivo*. In contrast, stability assays indicate that meprin α oligomers are less susceptible to proteolytic degradation and have improved thermal stability when compared to lower stoichiometry forms. The increased stability against proteolytic degradation seems to be meaningful considering the secretion of meprin α into compartments with high proteolytic potential such as the lumen of intestine and proximal kidney tubuli.

Lastly, it has been established that differential recognition and cleavage of substrates is dependent on enzyme localisation, such as membrane bound versus soluble meprin β. As is the case with TPPII, the formation of a helix could also function to sequester and concentrate activity to a localised region (67, 68). The effect of localised activity would not be captured in our *in vitro* assays, however *in vivo* it may be important for the catabolic function in the lumen of the intestine and proximal kidney tubuli, and additionally, for the controlled degradation of the extracellular matrix. Intriguingly, in some tissue contexts such as the epidermis, meprin α expression is observed to be specifically localised in particular subregions (e.g. the stratum basale). Owing to their size, meprin α filaments may have limited capacity to diffuse into neighbouring tissue areas thereby restricting the ECM remodelling activity to appropriate regions.

Considering these finding and our structures, we mapped frequently mutated residues that are associated with disease to the domains that control meprin α oligomerisation. This localisation suggests that these mutations may drive disease by destabilising the oligomeric form. For example, by the release of higher proportions of freely diffusing meprin α dimers that exert function in a dysregulated manner (Figure 5). The restricted helical arrangement further suggests that some activators, substrates, and inhibitors may have limited accessibility to the meprin α pro-peptide and active site. However, we were unable to detect any significant differences between meprin α oligomeric forms when assessing small molecule and globular substrates, nor small molecule and globular inhibitors. Indeed, structural studies of human fetuin-B in complex with meprin α are consistent with crystal structures of murine fetuin-B and meprin β (96). Unlike meprin β, these inhibitory complexes were observed to form supramolecular helical packing as a result of the intercalated arrangement of meprin α and numerous fetuin-B molecules.

**Fig. 5.**
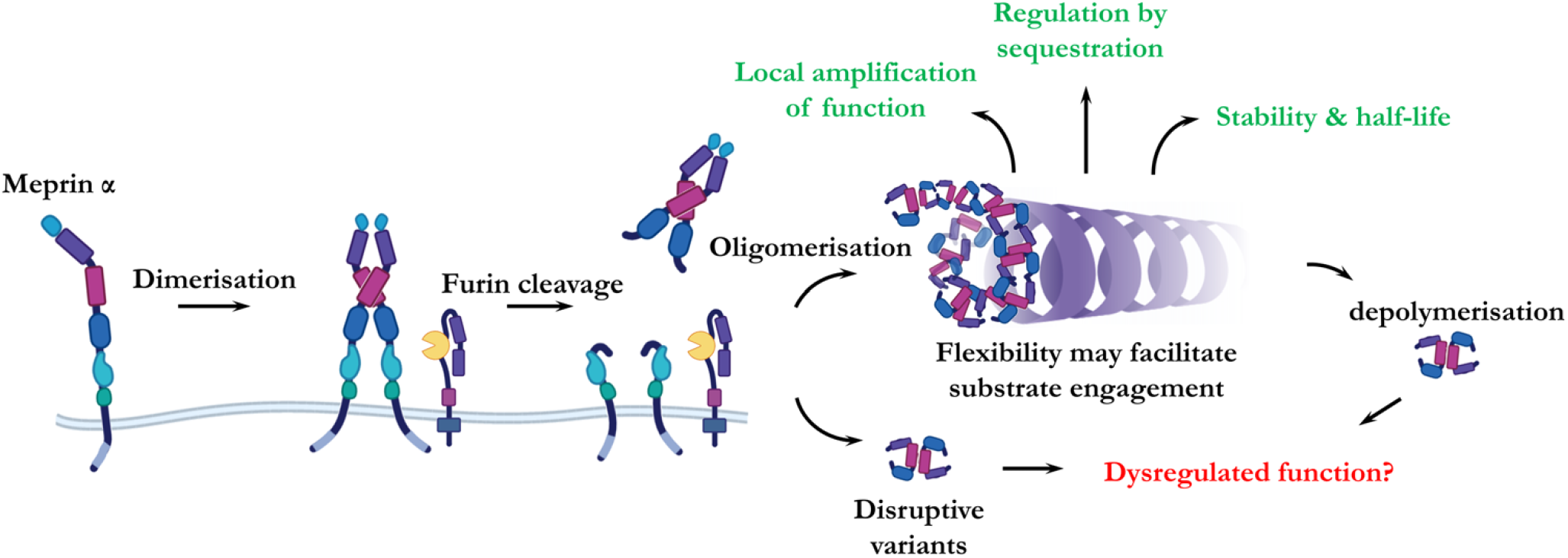
Proposed mechanism of action of meprin α helices. Monomeric full length meprin α dimerises during expression and is proteolytically shed from the membrane bilayer via furin-protease activity. Shed meprin α homodimers oligomerise via interactions of MAM/MATH domains forming large oligomeric helices. These helices function to locally amplify proteolytic activity as well as regulate the extent of activity by sequestrating the protease to specific cellular and tissue locations. Further, oligomerisation putatively improves *in vivo* stability and half-life. Disruptive mutations that interfere with helix formation or increase rate of depolymerisation are postulated to drive dysregulated and detrimental meprin α function.

Taken together, our findings suggest that differences between stoichiometric forms of meprin α may depend entirely on their localisation which, consequently, dictates their distinct activity profiles (15, 48). Ultimately the impact of these effects on biological function, if any, remains to be shown. It is currently unclear what role meprin α oligomerisation mediates at a cellular level. Previous studies suggest the tissue and cellular localisation of meprins underpins both correct function and regulates activities of these key proteases (15, 48). Therefore, we suggest meprin α oligomerisation may be a regulatory mechanism that functions to either stabilise, sequester and/or locally amplify proteolytic activity (Figure 5). Further studies in these regards will be informative in understanding the role of meprin α in cancers and other human diseases. For example, studies *in vivo* to assess the physiological effects of meprin α depolymerisation and the importance of meprin α oligomerisation in normal ECM homeostasis.

## Material and methods

### Expression, purification, activation

Expression of pro-meprin α (and variants) was achieved in Schneider-2 Drosophila cells (S2 cells). Briefly, the sequence of pro-meprin α (V22-S600; uniprot Q16819) containing an N-terminal Strep-Tag was cloned into pMT/BiP/V5, enabling stable cell lines to be produced. The pro-meprin α variants C308A, R372T and R372A were produced by site-directed mutagenesis. To induce the production of pro-meprin α, S2 cells were grown in Schneider’s Drosophila Medium (Biowest) supplemented with 1 mM copper sulphate and 0.05% PluronicTM F-68 at 28 °C and 80 rpm for two days. The supernatant was harvested by centrifugation and immediately purified by hydrophobic interaction chromatography applying expanded bed adsorption (HIC-EBA, equilibration buffer 30 mM Tris-HCl pH 7.4, 1.5 M ammonium sulphate; elution buffer 30 mM Tris-HCl pH 7.4, 100 mM NaCl). The eluate of HIC-EBA was subjected to affinity chromatography using Strep-Tactin^®^ column (5 ml cartridge, GE Healthcare Life Science; equilibration buffer 30 mM Tris-HCl pH 7.4, 100 mM NaCl; elution buffer 30 mM Tris-HCl pH 7.4, 100 mM NaCl, 2.5 mM desthiobiotin). Finally, pro-meprin α was activated by trypsin cleavage applying immobilized trypsin on magnetic beads (PT3957-1, Takara, buffer 30 mM Tris-HCl pH 7.4, 100 mM NaCl, 5 mM CaCl_2_).

### Animal work

Mouse urine was collected in metabolic cages for 24 h. The animal experiment was approved by the responsible animal ethics committee of the state of Saxony-Anhalt, Germany (Landesverwaltungsamt Sachsen-Anhalt, Department of Consumer Protection and Veterinary Affairs, Halle (Saale), Saxony-Anhalt, Germany) under the following approval number: 42502-2-1473 MLU.

### Kinetic analysis

The determination of enzymatic activity was based on the cleavage of the fluorescent peptide substrate Abz-YVADAPK(Dnp)G-OH. Measurement of kinetic parameters was performed in 384 well black plates in a volume of 60 μl (assay buffer 50 mM HEPES, 150 mM NaCl, pH 7.4, 0.05% Brij). Enzyme solution in buffer was applied and subsequently substituted with inhibitor/DMSO normalisation (Digital Dispenser D330e, Tecan, Switzerland) and preincubated at 30 °C for 20 min. After addition of substrate reaction was measured at excitation/emission wavelength 340/420 nm on a plate reader (Clariostar, BMG Labtech, Germany). For estimation of first order rate constants of kcat/Km substrate concentrations from 0.8 to 83 μM were used. Concentration of active enzyme was determined to be 3–5 nM by titration of the active site using a tight binding inhibitor. The IC50 were estimated at a substrate concentration of 10 μM varying inhibitor from 10 μM to 5 nM (compound 10d) and 200 nM to 0.03 nM (fetuin-B). Kinetic parameters were determined at least in triplicates in separate experiments and evaluated with GraphPad Prism software (Graphpad Software, Inc., USA). Mouse fetuin-B was a generous gift from Hagen Körschgen and Walter Stöcker (University Mainz).

### Turnover of protein substrates

Different enzymesubstrate ratios of meprin α wild type and meprin α C308A were incubated with human tropoelastin, human fibronectin and rat collagen I. In the case of tropoelastin (kindly gifted by Mathias Mende, group of Prof. Pietzsch, Martin-Luther University Halle-Wittenberg), the reaction was carried out in 50 mM Tris-HCl pH 7.4 buffer at 37 °C and molar ratios of enzyme to substrate of 1:10^4^ and 1:10^5^. Samples (10 μg) were removed every 30 min for of 3 h. The cleavage of fibronectin (ab209886, Abcam) was assessed in 50 mM HEPES pH 7.4 buffer containing 150 mM NaCl. Enzyme to substrate ratios of 1:10^4^ and 1:10^5^ were tested. Reaction conditions were as described above. Collagen I from rat tail was prepared according to Gorisse *et al* (69) and the reaction was carried out in 50 mM HEPES pH 7.4 at 37 °C at a ratio of 1:10^4^. All cleavage products were evaluated by reducing SDS-PAGE, followed by Coomassie-staining.

### Proteolytic stability

A total of 9 μl of enzyme (2 μM) were supplemented with 1 μl Trypsin/Plasmin (2 μM) or buffer and incubated at room temperature. After 20 min, reactions were stopped by addition of 2.5 μl 5-fold reducing sample buffer. Samples were heated and subjected to analysis by 10% SDS-PAGE.

### NanoDSF

NanoDSF measurements were conducted on a Prometheus NT.48 (NanoTemper) instrument. Each meprin α variant was standardised to a protein concentration of 0.1–0.3 mg ml^−1^ in 30 mM Tris-HCl, 100 mM NaCl, pH 7.4. Thermal unfolding was monitored by the intrinsic tryptophan fluorescence at emission wavelengths 350/330 nm in 1 °C min^−1^ increments from 20 °C to 95 °C. The apparent melting temperature (T_m_) was determined as the maximum of the first derivative of the 350/330 nm ratio. Each variant was measured in three independent experiments.

### MADLS/SEC-MALS

Multiangle dynamic light scattering analyses of meprin α was performed using a Zetasizer Ultra (Malvern Panalytical Ltd., UK) and He-Ne laser at 632.8 nm and constant power of 10 mW at 30 °C. Concentration of protein was 0.3 mg ml^−1^ in 30 mM Tris-HCl, 100 mM NaCl, pH 7.4. Detector angle for polydispersity index (PI) was 173°. The results are presented as the average value of three to five experiments. For size exclusion-multiangle dynamic light scattering, meprin α C308A (100 μl at 0.74 mg ml^−1^) and meprin α R372T (100 μl at 0.44 mg ml^−1^) were applied onto an AdvanceBio SEC column (300 Å, 7.8×300 mm, Agilent Technologies). The data was evaluated using ASTRA^®^ 6 (Wyatt Technology).

### MALDI-TOF-MS

For MALDI-TOF mass spectrometry all samples were purified using C4 ZipTip^®^ pipette tips (Merck) before analysis as described in the supplied manual. The purified sample (1 μl) was mixed with 1 μl of DHAP-matrix (7 mg 2,5-dihydroxyacetophenone, 375 μl ethanol, 125 μl of 16 mg ml^−1^ di-ammonium hydrogen citrate) and 1 μl of 0.1% TFA and applied onto a metal target plate. The samples were analyzed in linear positive mode (LP_30-210 kDa according to Bruker Daltonics) using an AutoflexTM speed MALDI-TOF/TOF device (Bruker Daltonics). Protein Calibration Standard I and II (Bruker Daltonics) were applied for calibration of the device.

### Cryo-EM and cryo-ET sample preparation and data collection

For tomography samples, Quantifoil Cu R 2/2 grids were glow-discharged for 30 s in a Pelco EasyGlow, and 3 μL of meprin α (0.5 mg ml^−1^ in 30 mM Tris-HCl buffer, 100 mM NaCl, pH 7.4) was mixed with 10 nm gold nanoparticles and was applied to the glow discharged surface and blotted at 4 °C and 100% relative humidity for 3 s and a blot force of −3 using the Vitrobot IV System (Thermo Fisher Scientific). Grids were plunged in 100% liquid ethane. The grids were stored under liquid nitrogen until TEM data collection. Tilt series were acquired at 300 kV on a FEI Titan Krios G1 and digitised on a postGIF K2 Summit Direct Electron Detector. A dose-symmetric tilt acquisition scheme was used with 3° increments, an electron dose of 2.6 e^−^ A^−2^ per tilt. Micrographs were acquired in dose fractionation mode with 0.5 e^−^ A^−2^ frame^−1^ at a pixel size of 0.186 nm × 0.186 nm.

Similarly for single-particle cryo-EM, initial grid freezing conditions were tested and screened on a Tecnai T12 electron microscope (Thermo Fisher Scientific). Blotting was carried out as above, with the following modifications, a blot time of 2.5 s, blot force of −5 and drain time of 1 s were used. The grids were stored under liquid nitrogen until TEM data collection. Briefly, dose fractionated movies were collected on a Titan Krios (Thermo Fisher Scientific), equipped with a Quantum energy filter (Gatan) and Summit K2 (Gatan) or K3 (Gatan) direct electron detector. Data acquisition was performed using either SerialEM (70) or EPU (Thermo Fisher Scientific). Meprin α/fetuin-B complex was made immediately prior to freezing, by mixing stoichiometric quantities (final concentration 1 mg ml^−1^) and allowing to incubate for 3 minutes at room temperature. His-tagged recombinant human fetuin-B (11834-H08H) was purchased from SinoBiological. Similarly, compound 10d was added in excess (~8 fold Ki) and allowed to incubate for 1 minute, prior to freezing.

### Cryo-EM and cryo-ET image analysis

Dose fractionated movies were firstly compressed to LZW TIFF format with IMOD (71) to save disk space. Correction of beam induced motion and radiation damage was performed with MotionCor2 (v1.3.0) (72). Corrected frames were dose-weighted and averaged for all further processing. Tilt series were aligned and backprojected using IMOD (71, 73, 74), and CTF corrected using NovaCTF (75). Subtomogram averaging was performed using Dynamo (76). Subtomograms were extracted following a cylindrical geometry and the asymmetric unit used for alignment corresponded to 2 protein dimers. Subtomogram averages were later used as initial volumes for single particle analysis.

Single-particle data was analysed as follows. Initial estimates of CTF were performed in CTFFIND (4.1.13) (77) or Warp (1.0.7) (78). A combination of manual and automated filament particle picking was performed by hand or with crYOLO (1.8.1) (79, 80) operating in filament mode. A small, hand-picked subset of images were used to train crY-OLO. Particles were extracted within 400-pixel boxes and normalised within RELION (2.1-3.1) (81–83). Initial rounds of 2D classification in cryoSPARC (84, 85) and RELION were used to discard malformed particles and poor-quality images. Asymmetric refinements were performed in RELION. Flexibility resulted in large-scale, continuous conformational heterogeneity between the images. This was apparent in 2D class averages which displayed clear secondary features at the centre of the averages, with diffuse signal and poor coherency that worsened moving away from the centre along the helix lengthwise. Three-dimensional variability analysis in cryoSPARC revealed numerous modes of flexibility (86). This also prevented symmetry determination by layer line analysis. Resultantly, all the refinements of the full helical assembly that were attempted, gave rise to severely limited reconstructions with resolution on the order of nanometres.

The best reconstructions of helical complexes were used to guide the placement of meprin β dimer modes (PDB-4GWN) (53). A mask of approximately two dimers was used to carve ~6 regions of interest from the reconstruction. A small space was left between regions as a buffer to accommodate some variation in the placement of models. These regions of interest were individually removed from the reconstruction, to generate masks of the remainder of the helix. These masks were subsequently used for partial signal subtraction and localised particle extraction within a modified version of *localised_reconstruction.py* (87) or directly within RELION.

A carved segment of the original helix was re-boxed and used an initial volume. Localised reconstruction of these portions of helix gave rise to a notably more homogeneous volume that displayed secondary structures. Subsequent 2D and 3D classifications of these segments were performed in RELION or by heterogeneous refinement in cryoSPARC. Homogeneous refinements in cryoSPARC were performed to obtain optimised alignment and offset parameters. Peripheral regions of the localised reconstruction corresponding to the boundaries of the subtracted regions were noisy and diffuse. Therefore, these areas were subtracted and further rounds of local refinement in cryoSPARC were performed. CTF parameters were refined and corrected in cryoSPARC or relion, including anisotropic magnification, beam tilt, trefoil, tetrafoil and astigmatism (83). Finally, non-uniform refinement was employed to account for variation in resolution across the reconstruction that gave rise to minor errors in alignments (85).

For reconstructions of fetuin-B/meprin α, RELION was unable to solve the global alignment problem. Therefore, asymmetric reconstructions of the whole helix were performed in cryoSPARC. Localised sub-particle extraction was performed in RELION and global alignment of the extracted subregions was performed in RELION (here cryoSPARC failed to solve the global alignment problem). Finally, local non-uniform refinement in cryoSPARC with gaussian prior yielded the final reconstruction. These final particles and optimised metadata were exported from cryoSPARC for 3D classification within RELION. Low resolution signal dominated and therefore the CTF was ignored until the first zero giving rise to markedly improved classification.

For all maps local resolution was estimated in RELION with a windowed FSC. Map sharpening was performed in cryoSPARC, RELION or deepEMhancer (88). Various amplitude corrected maps were employed for model building. Conversion between cryoSPARC and RELION were performed with *pyem* (89).

### Model building

An initial model of full length pro-meprin α were generated via homology with meprin β (PDB-4GWN) (53) using the SWISS-MODEL server (90) and rigid body fit into the cryo-EM reconstruction with UCSF Chimera (v1.13) (91). Subsequently, the model was flexibly fit and interactively refined in ISOLDE (92), ChimeraX (v1.3) (93) and Coot (94, 95). Where possible, N-linked glycosylations were modelled in Coot. Models of the active form and inhibitor-bound form were generated similarly, using the model of pro-meprin α as a starting template. Lastly, molecular docking analysis was carried out using the fetuin-B/meprin β crystal structure (PDB-7AUW) (96) superimposed onto the meprin α helix. Subsequently, a model of the meprin α/fetuin-B complex was generated from the model of active meprin α and the AlphaFold model of human fetuin-B (97, 98) (AF-Q9UGM5 v2). These were rigid body fit into the cryo-EM reconstruction and were subsequently flexibly fit and interactively refined in ISOLDE/ChimeraX using secondary structure and all atom constraints to prevent the model from diverging. All models were subject to real space refinement and validation in PHENIX (99, 100) and MolProbity (101).

## DATA AVAILABILITY

Cryo-EM maps are deposited in the Electron Microscopy Data Bank (EMDB), while atomic coordinates are available from the RCSB Protein Data Bank under the following accession codes. Pro-meprin α (tetramer), EMD-26419 and PDB-7UAB. Pro-meprin α (single subunit), EMD-26420 and PDB-7UAC. Full meprin α helix in the active state (C1 reconstruction), EMD-26421. Meprin α in the active state (single subunit), EMD-26422 and PDB-7UAE. Meprin α in complex with small molecule inhibitor (tetramer), EMD-26423 and PDB-7UAF Meprin α in complex with the native fetuin-B inhibitor (tetramer), EMD-26426 and PDB-7UAI. Full meprin α helix in complex with the native fetuin-B inhibitor (C1 reconstruction), EMD-26424. Additional information and data are made available upon reasonable request from the corresponding author.

## ACKNOWLEDGEMENTS

CBJ acknowledges the support of the Australian Government RTP scholarship. We thank the staff of the Monash Ramaciotti Centre for Electron Microscopy, the support of the MASSIVE supercomputer team and the Monash Macromolecular Crystallisation Facility. This project was supported by BMBF (APRA program), Germany. We gratefully acknowledge the organisational support by Vivoryon N.V. and in particularly the valuable scientific discourse and proofreading by Stefan Tasler. This preprint was formatted in LATEX using an adaptation of Ricardo Henriques’ template.

## AUTHOR CONTRIBUTIONS

D.S., C.F., M.W. expressed, purified, and analysed protein. D.S., C.F., C.J.L. prepared and screened cryo-electron microscopy samples. C.B.J, C.J.L., D.S. and C. F. processed and analysed cryo-electron microscopy data. C.B.J., C.J.L., D.S. and C.F. built and analysed protein models. C.B.J. wrote the first draft. C.B.J., D. S., C.F., C.J.L., J.C.W. co-wrote, revised, and edited the manuscript. C.B.J., D.S., C.J.L., produced figures. H.V., A.dM., C.J.L. collected cryo-electron microscopy data. D.R. and C.J. performed inhibitor docking and structural analysis. C.B.J, D.S. and C.F. produced figures. D.S., S.S., and J.C.W. acquired funding and provided supervision. All authors contributed to the discussion and interpretation of results.

---

**Fig. S1.**
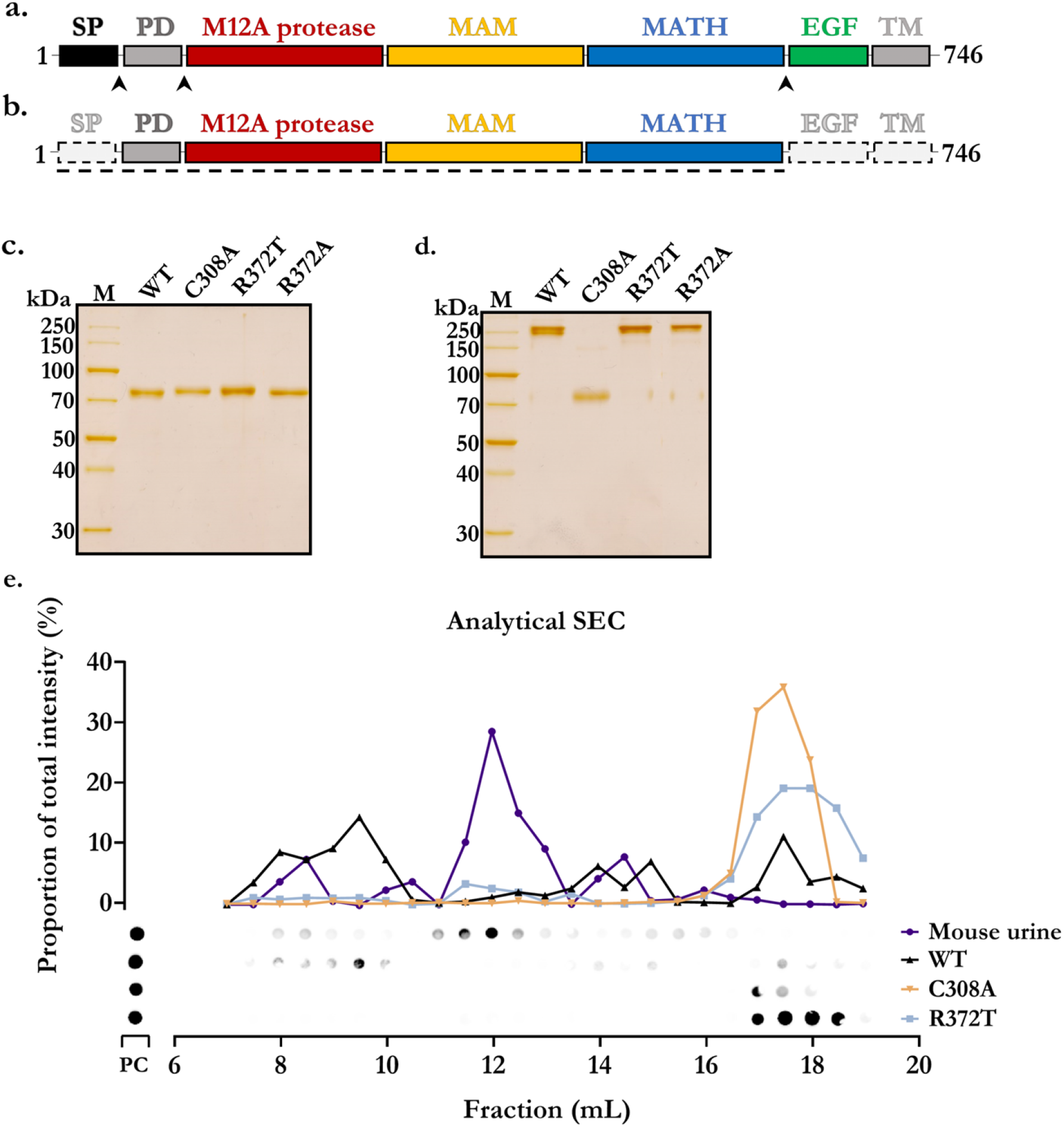
Meprin α expression and purification. **a.** Domain schematic of native meprin α consisting of a signal peptide (SP), pro-domain (PD), proteolytic (M12A protease) domain, meprin, A-5 protein, and receptor protein-tyrosine phosphatase μ (MAM) domain, TNF receptor-associated factor (TRAF or MATH) domain, epidermal growth factor-like (EGF) domain and a transmembrane region (TM). Arrows indicate sites of proteolysis during normal meprin α maturation. **b.** Recombinant construct employed herein. Dashed boxes represent regions that are not genetically encoded in the recombinant construct. **c.** Reducing 10% SDS-PAGE analysis of purified recombinant meprin α and variants. **d.** As in (b) with non-reducing conditions. **e.** Comparison of retention volume between recombinant meprin α (wild type, C308A and R372T) to native source meprin α isolated from mouse urine. Large oligomeric species were observed from the mouse urine after cisplatin treatment. These were, however, smaller than recombinant oligomeric meprin α.

**Fig. S2.**
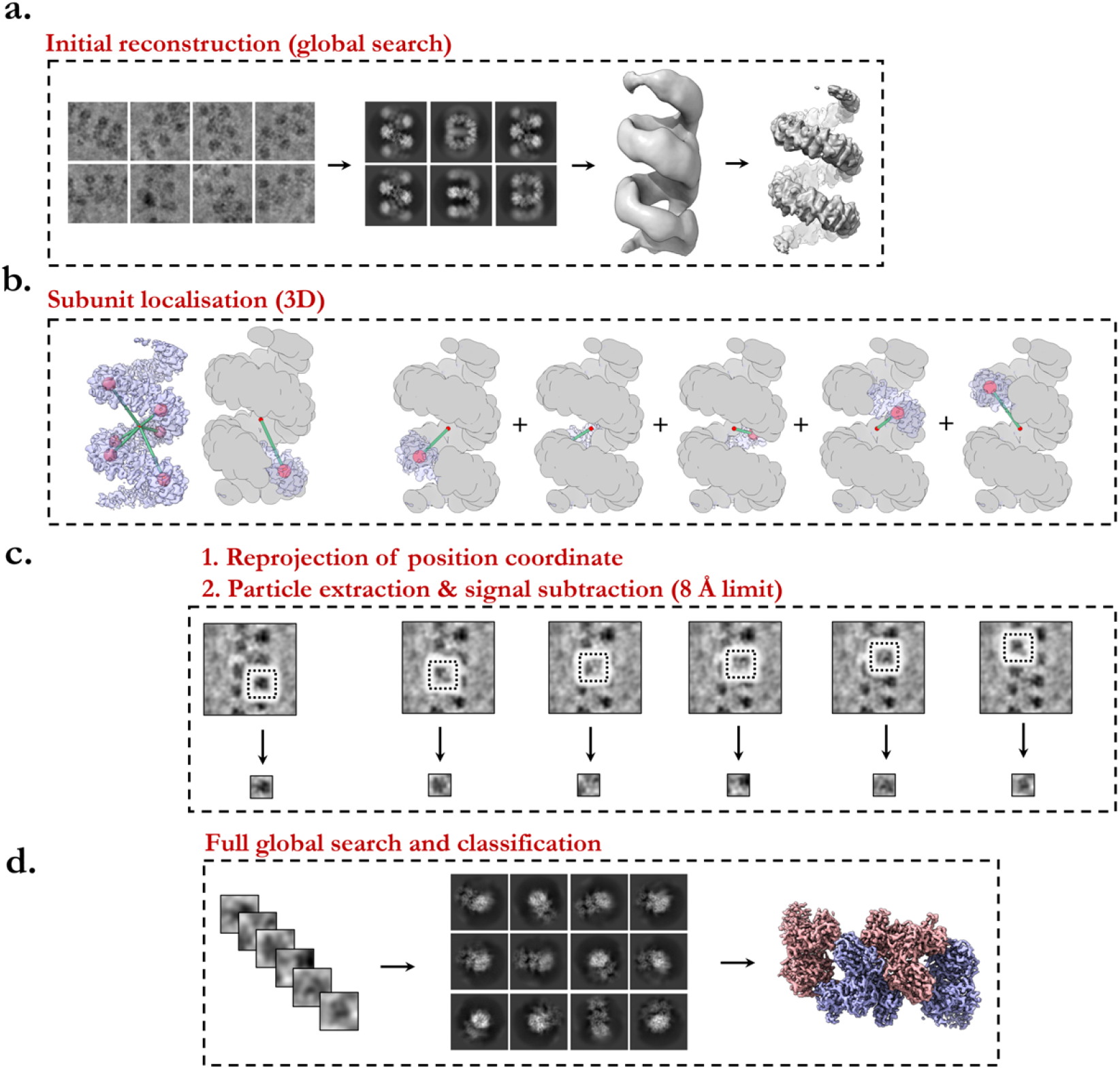
Summary diagram of cryo-EM image analysis strategy and reconstruction method. **a.** Helical segments are broken into 20-30 nm fragments and classified independently. An initial volume was created by sub-tomogram averaging. Refinement of a full helix was performed by single-particle analysis. **b.** Coordinates of meprin α tetramers were defined based on the final reconstruction of the full helix and used to reproject 2D coordinates onto the original extracted particles. **c.** Sub-regions of meprin α tetramers were re-extracted from the original particles within a smaller area and subsequently treated as independent single particles. d. Assumption of single particle behaviour enabled reconstruction of a tetramer localised reconstruction by performing global searches with C1 “symmetry”.

**Fig. S3.**
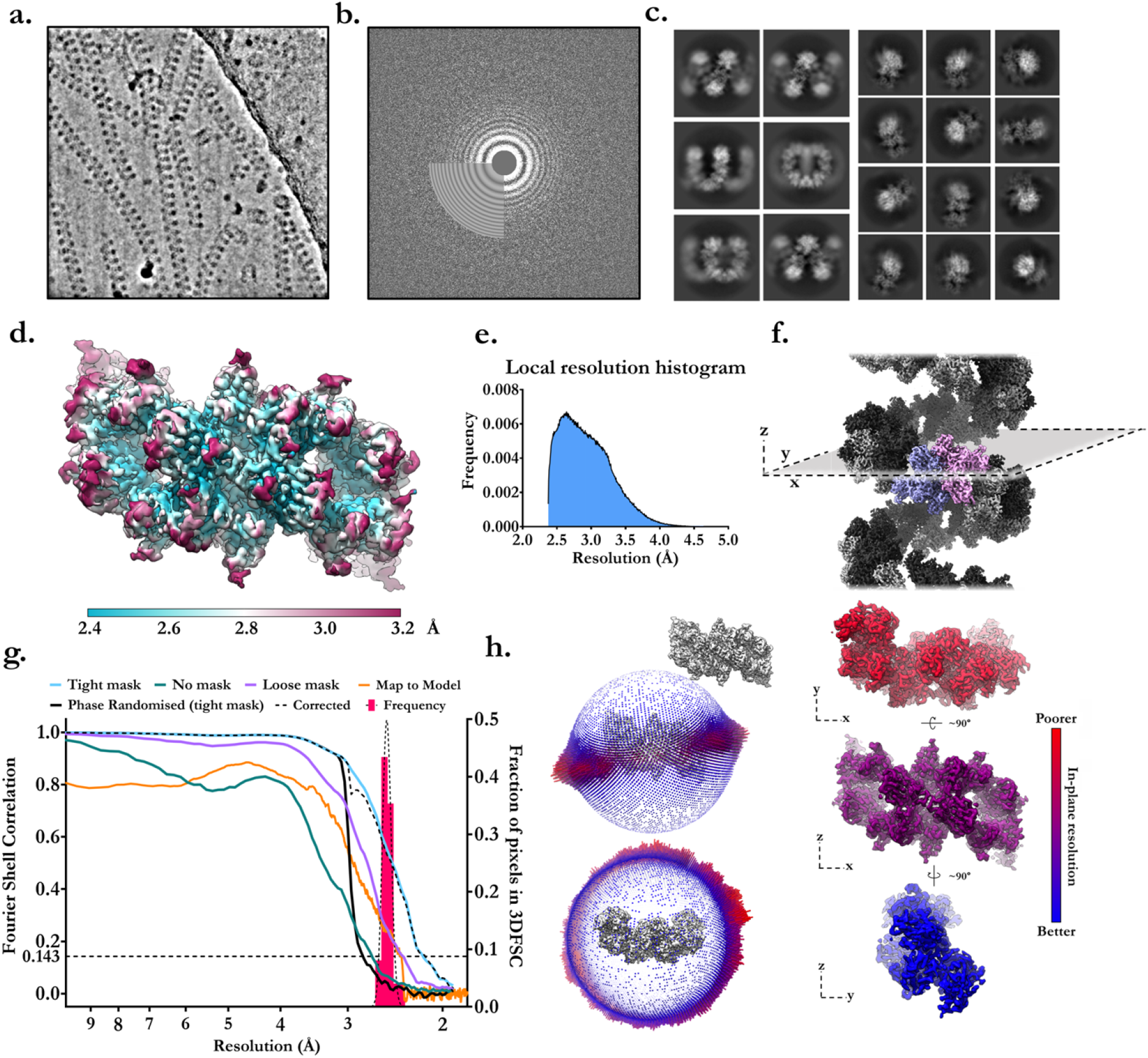
Summary figure of cryo-EM key statistics and analysis outcomes (compound 10d map). **a.** Representative micrograph of meprin α helices and, **b,** corresponding Fourier transform with high quality Thon rings. **c.** Class averages in 2D of meprin α helices and of meprin α tetramer subparticles after localised extraction. **d.** The final meprin α tetramer reconstruction coloured by local resolution. **e.** Corresponding per-voxel resolution frequency distribution. **f.** Composite map of the meprin α helix and corresponding directional resolution analysis. Maximum anisotropy is observed from the z-direction due to the tendency of helices to lie flat within the ice. **g.** Fourier shell correlations including tight, loose and no mask curves, phase randomised half-maps, mask corrected FSC and histogram of directional 3DFSC voxels (frequency). **h.** Angular distribution and orientation assignment of observed particles after refinement.

**Fig. S4.**
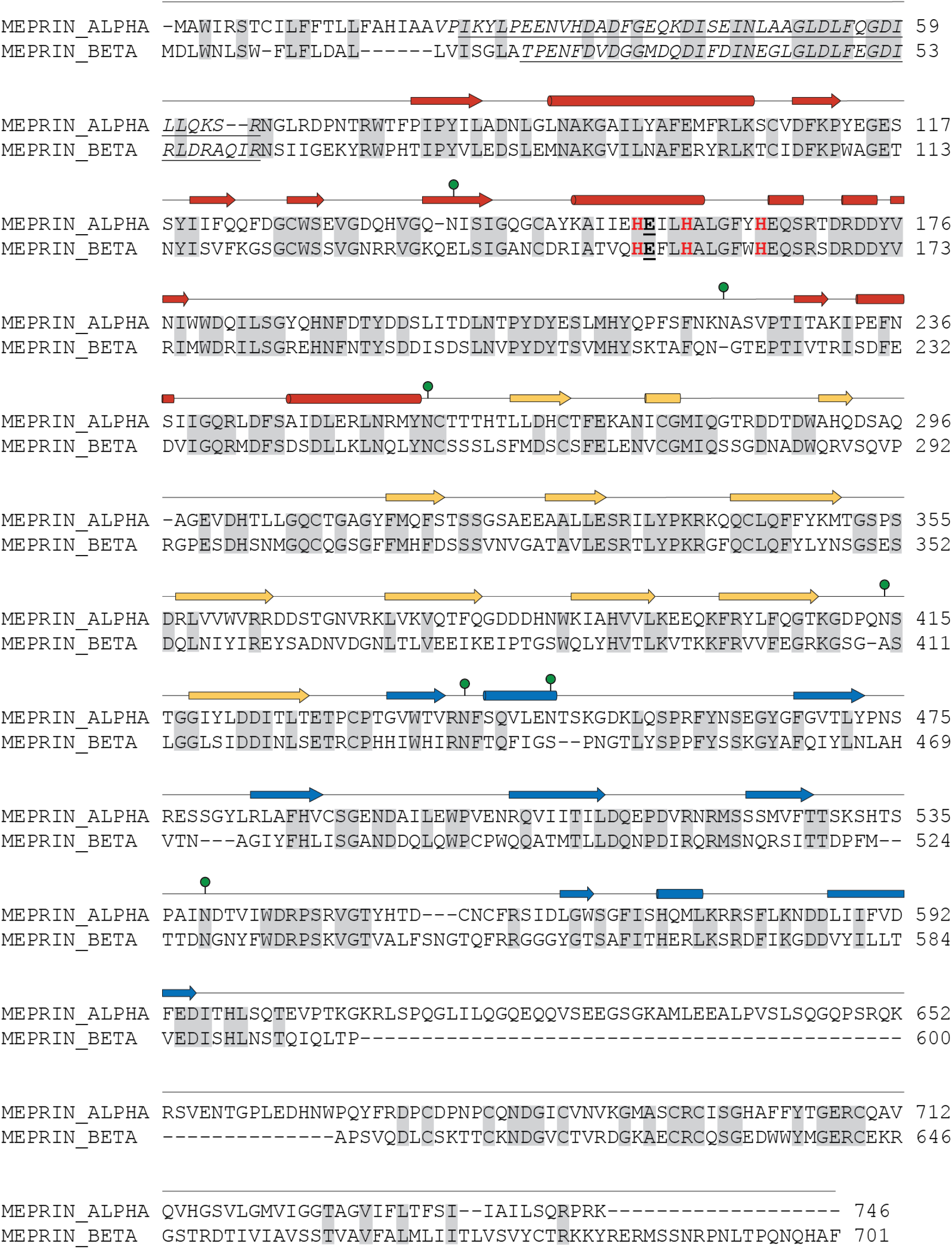
Sequence alignment of meprin α and meprin β. Secondary structure elements are shown above based on the structure reported herein, arrows represent β-strands and tubes represent α-helices. Green circles represent N-linked glycosylations. The active site triad of histidine residues are marked in bold font and red. The catalytic glutamic acid is black bold font emphasised.

**Fig. S5.**
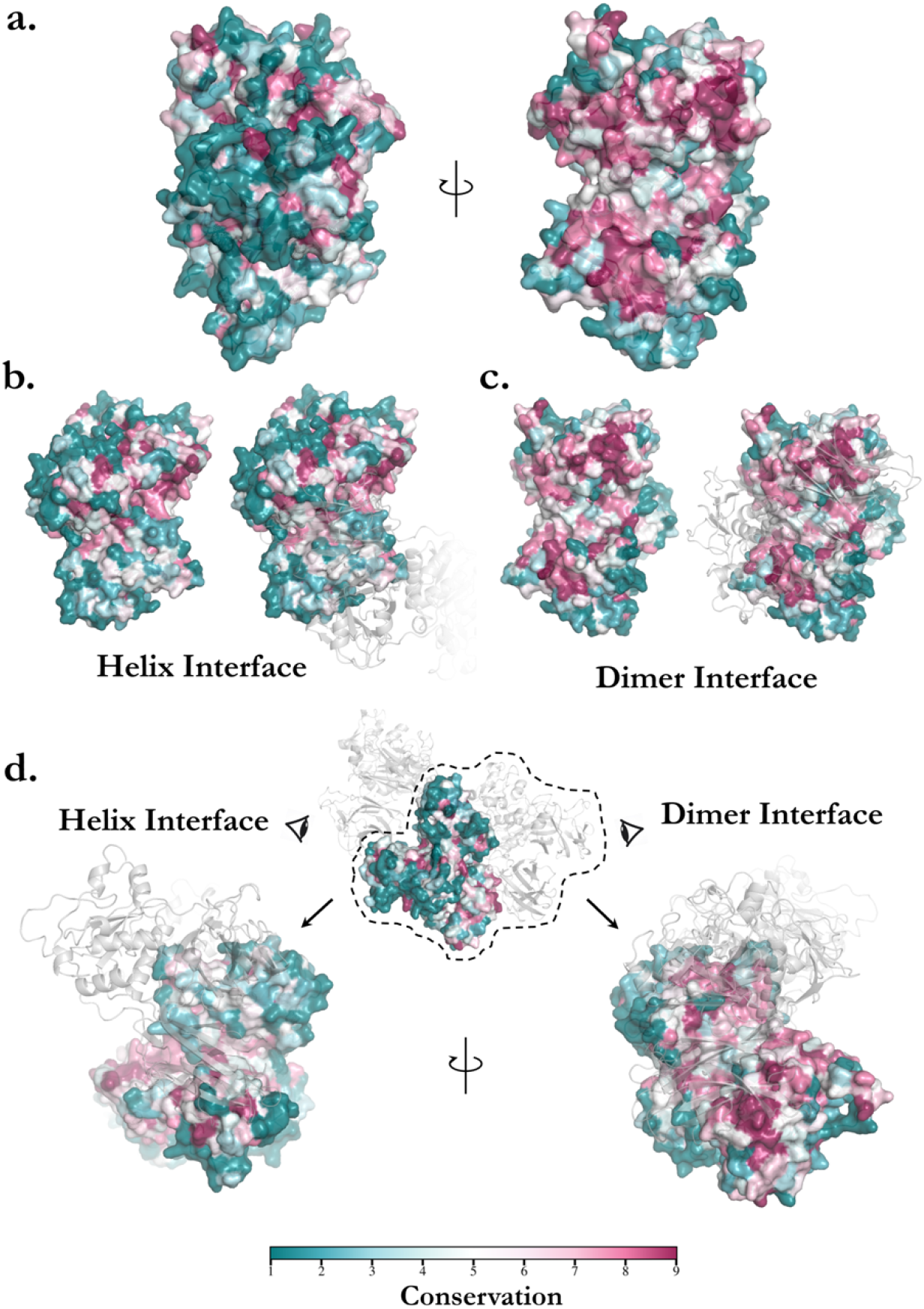
ConSurf analysis of meprin α dimer and oligomer interfaces. **a.** Monomer of meprin α with conserved surface rendering showing highly conserved dimeric interface (right). **b.** Focus of helical interface as conserved surface rendering (with adjacent monomer in transparent cartoon), showing conservation to some degree. **c.** Focus of dimer interface (with cartoon model of adjacent monomer in transparent), showing strong conservation corresponding to the binding site of the adjacent monomer. **d.** Single monomer of meprin α shown in context of the helical oligomer.

**Fig. S6.**
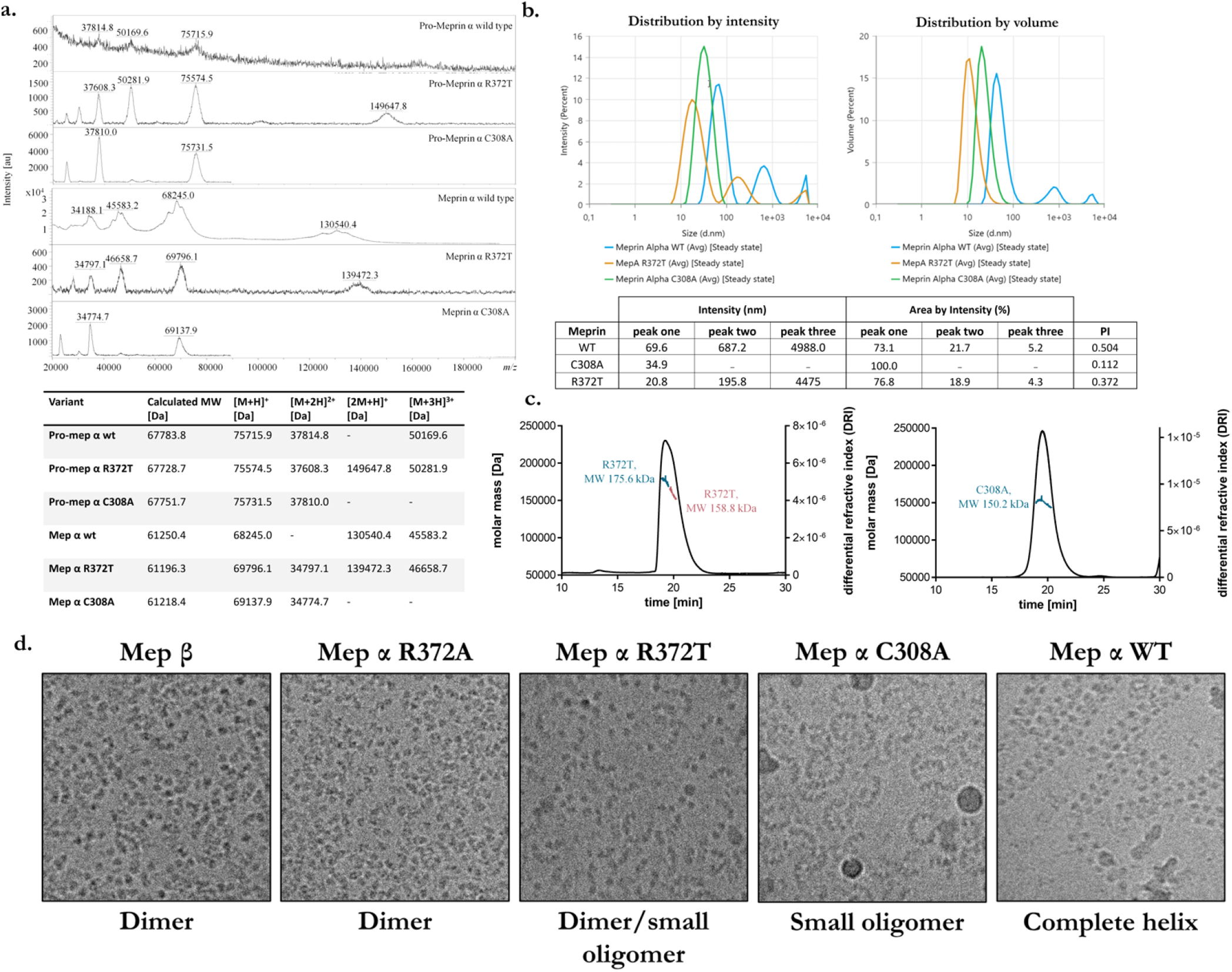
Biophysical characterisation of wild type meprin α and variants. **a.** MALDI-TOF analysis of wild type meprin α, C308A and R372T, as both zymogen and mature form. Peaks of about 75 kDa (zymogen) and 69 kDa (mature form) equal [M+H]^+^ or [M+2H]^+^, peaks of about 38 kDa (zymogen) and 35 kDa (mature form) equal [M+2H]^2+^. **b.** Comparative analysis of hydrodynamic radius by MALDS of meprin α and mutants C308A and R372T with the Zetasizer Ultra. Overlay of intensity particle size distribution (left). Overlay of volume particle size distributions results are presented as the average value of three to five experiments (right). **c.** SEC-MALS analysis of Pro-Meprin α variants C308A and R372T. Zymogenic meprin α C308A and R372T form dimeric structures, whereas for C308A a homogenous peak was observed, for R372T a heterogenous peak was detected, including two forms of dimeric pro-meprin α R372T. **d.** Cryo-TEM (120 kV) of meprin α and β. Mutant variants of meprin α form dimers and small oligomers.

**Fig. S7.**
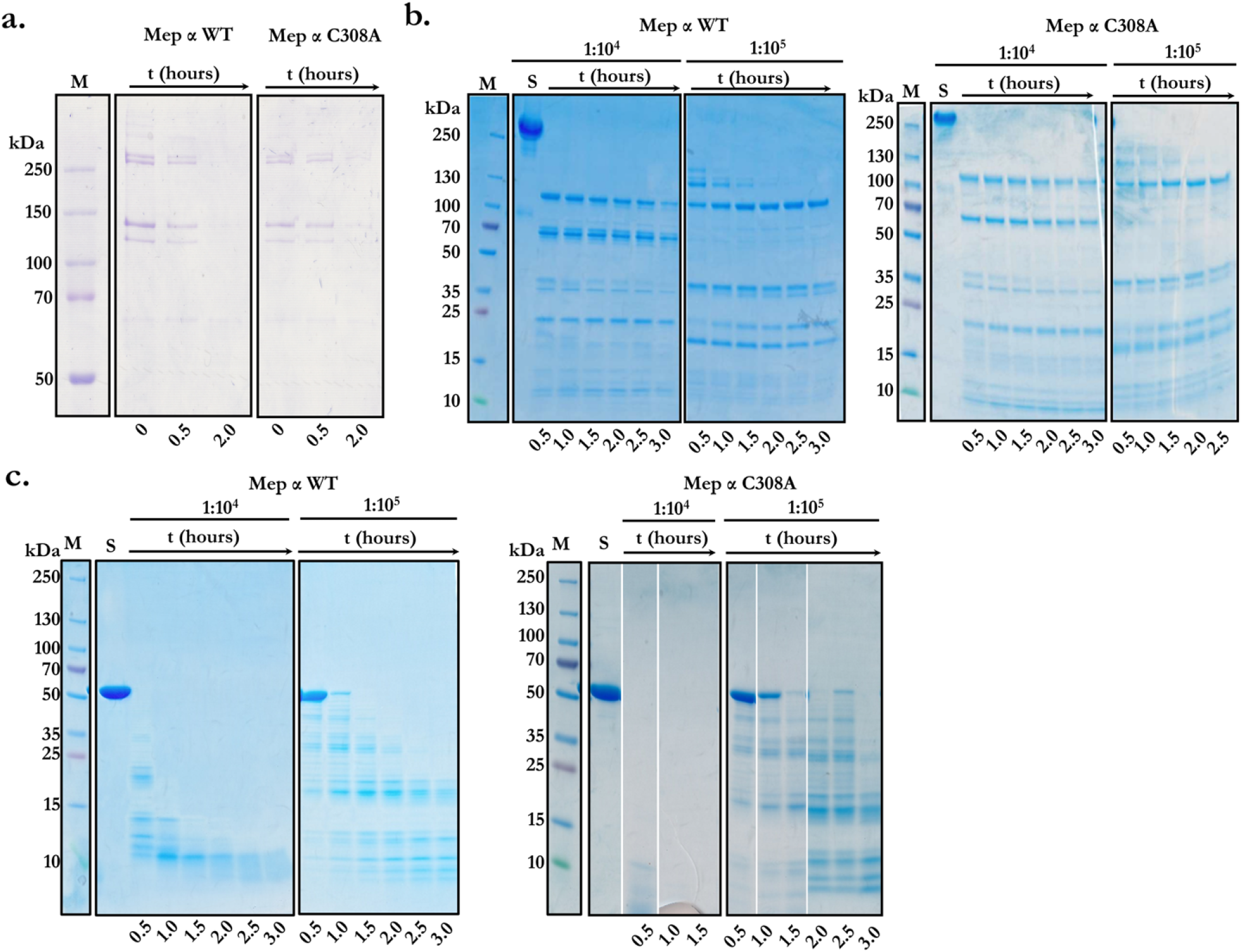
Meprin α activation, substrate degradation and activity. Oligomeric state does not appear to drastically affect substrate specificity or rate of degradation. **a.** Cleavage of rat tail procollagen by wild type meprin α and variant C308A. Specific cleavage of procollagen by both meprin α variants result in same cleavage pattern. Samples analysed using reducing 10% (w/v) SDS-PAGE, visualized by Coomassie-staining. **b.** Cleavage of human fibronectin by wild type meprin α and variant C308A. Specific cleavage of Fibronectin by both meprin α variants result in same cleavage pattern. **c.** Cleavage of human tropoelastin by wild type meprin α and variant C308A. Complete degradation of tropoelastin by wild type meprin α within 3 h (molar ratio of 1:10^4^) and by meprin α C308A within 30 min (molar ratio of 1:10^4^). Samples analysed using reducing SDS-PAGE (4-20% (w/v) gradient gel), visualized by Coomassie-staining. In each sample 10 μg of substrate were applied, as well as a control of substrate only (S; substrate incubated for 3 h at 37 °C in the absence of meprin α).

**Fig. S8.**
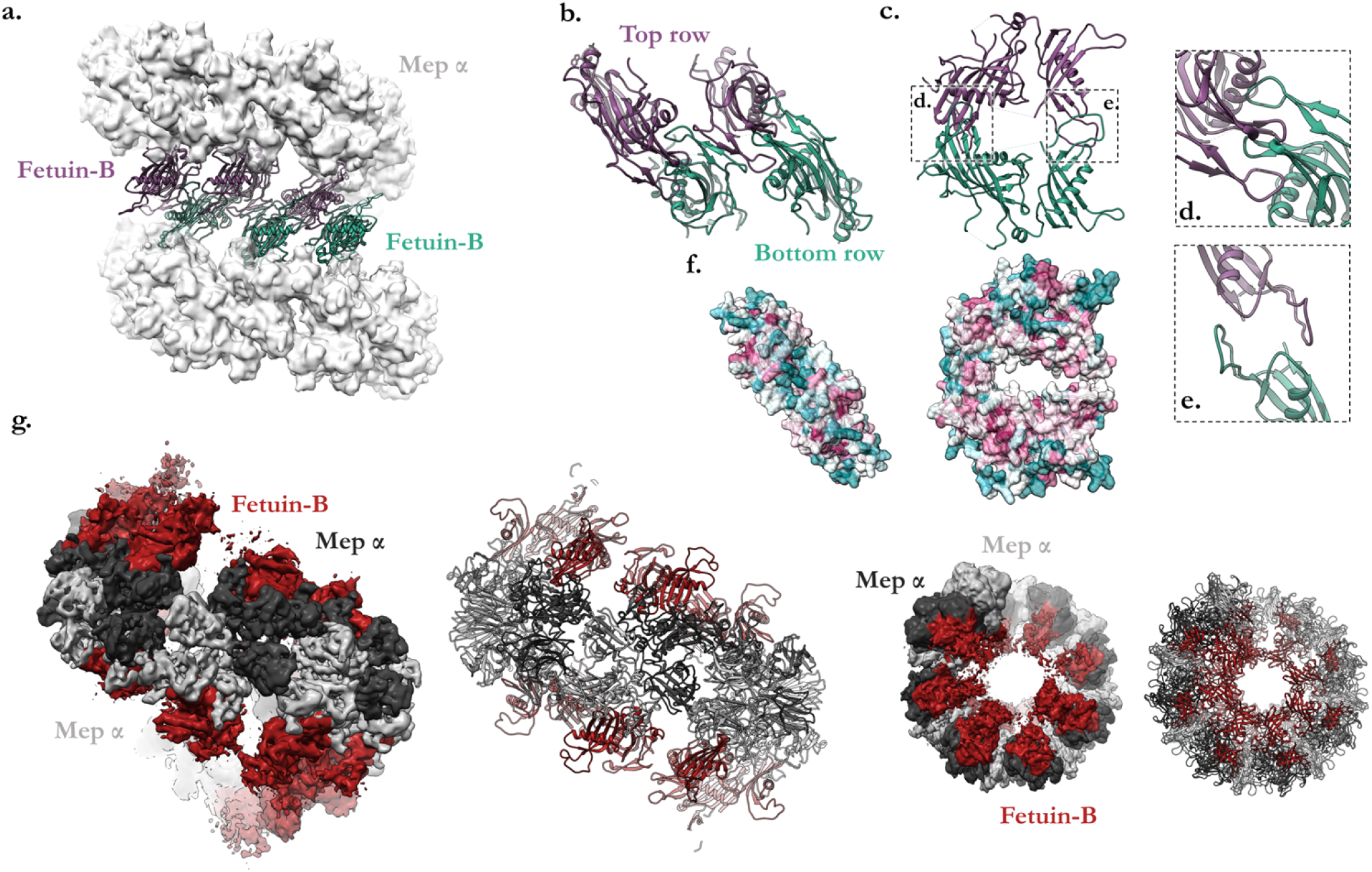
Molecular docking and 3.7 Å cryo-EM reconstruction of fetuin-B inhibitor bound to meprin α. **a.** Rigid body fit of murine fetuin-B/meprin β crystal structure (PDB 7AUW) to helical structure of meprin α. Arrangement of fetuin-B reveal monomers may pack into a slanted intercalated state that is not significantly prohibited by steric clashes. **b.** View of a tetramer of fetuin-B based on meprin α docking reveals potential interactions to form a higher-order inhibitory filamentous complex are possible. **c.** Side view of single fetuin-B dimer forms a “horseshoe” where putative interactions between inter-subunit fetuin-B domains may occur shown in (d), and (e). **d, e.** Models are not refined, rigid body fitting results in some minor clashes. **f.** The predicted oligomeric interface corresponds to an evolutionarily conserved interface revealed by ConSurf analysis. **g.** Side and top views of the cryo-EM reconstruction of human fetuin-B (red) in complex with meprin α (grey, black) at 3.7 Å resolution. Fetuin-B is observed to pack intimately within the meprin α active groove and intercalate as a secondary helix.

**Fig. S9.**
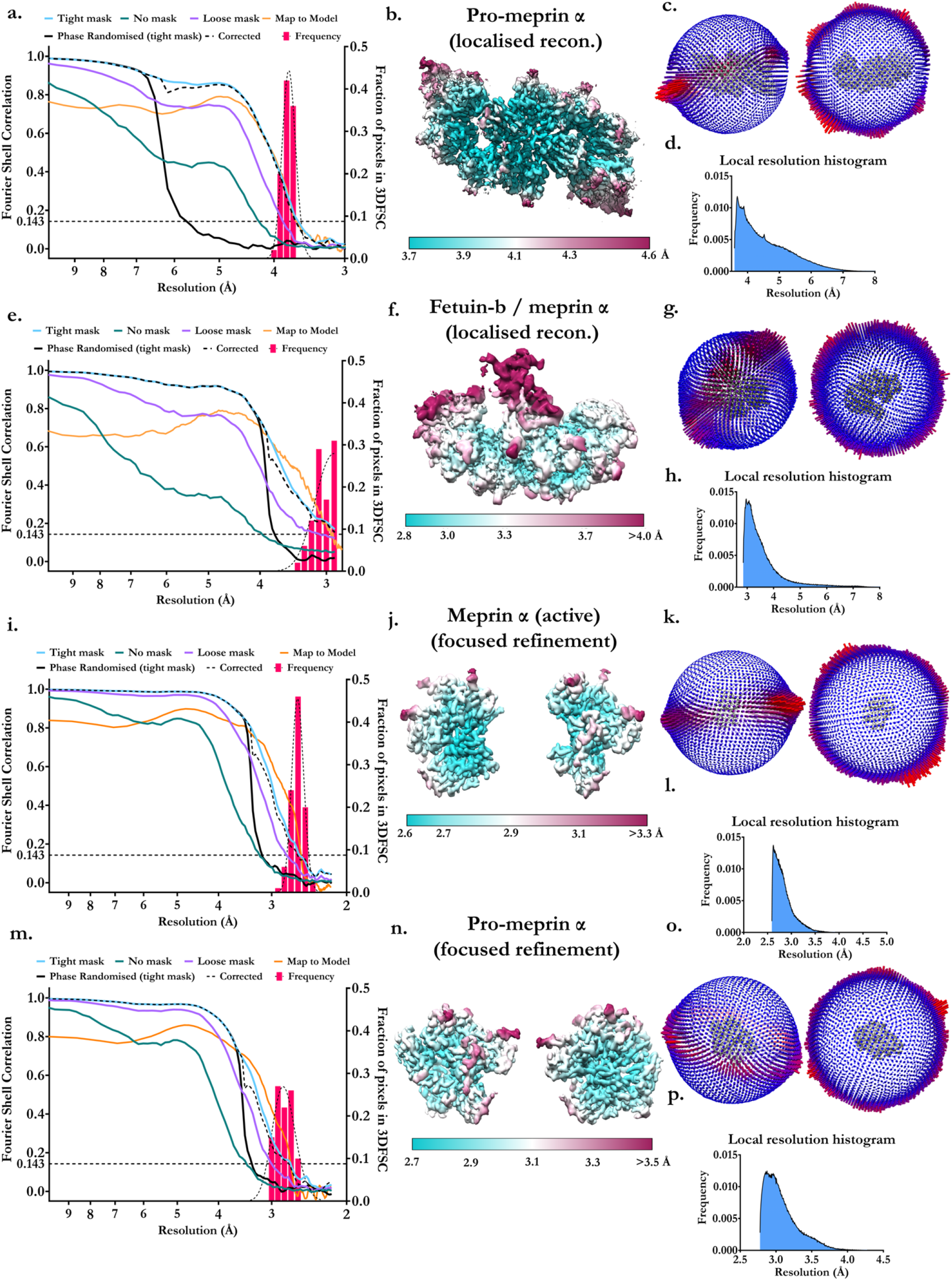
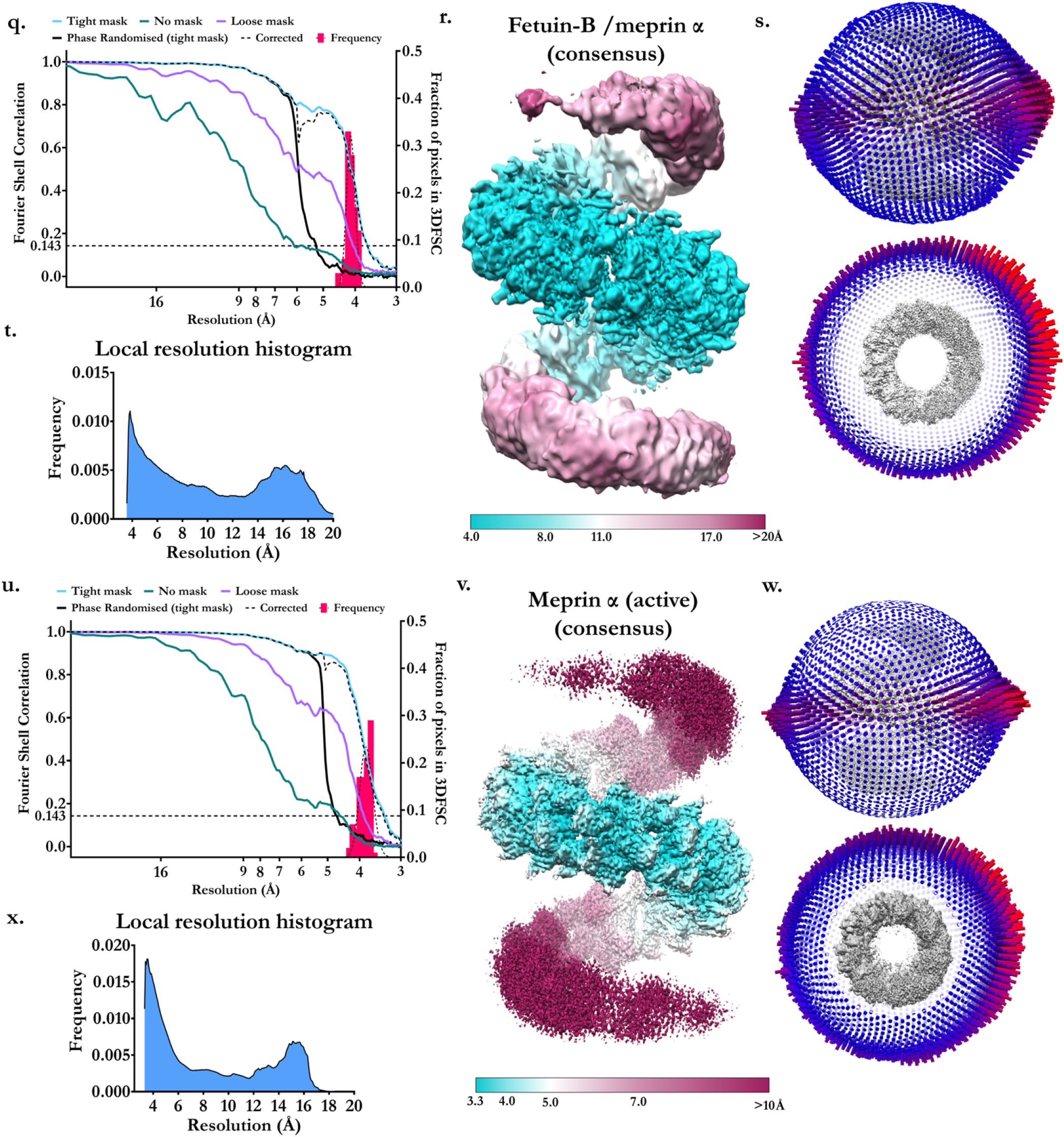
Summary figure of cryo-EM key statistics and analysis outcomes (zymogen, active and fetuin-B maps). **a., e., i., m.** Fourier shell correlations including tight, loose and no mask curves, phase randomised half-maps, mask corrected FSC, map-to-model FSC and histogram of directional 3DFSC voxels (frequency). **b., f., j., n.** Final reconstruction coloured by local resolution. **c., g., k., o.** Corresponding per-voxel resolution frequency distribution. **d., h., l., p.** Angular distribution and orientation assignment of observed particles after refinement. **q., u.,** Fourier shell correlations including tight, loose and no mask curves, phase randomised half-maps, mask corrected FSC and histogram of directional 3DFSC voxels (frequency). **r., v.** Final reconstruction coloured by local resolution. **t., x.** Corresponding per-voxel resolution frequency distribution. **s., w.** Angular distribution and orientation assignment of observed particles after refinement.

**Fig. S10.**
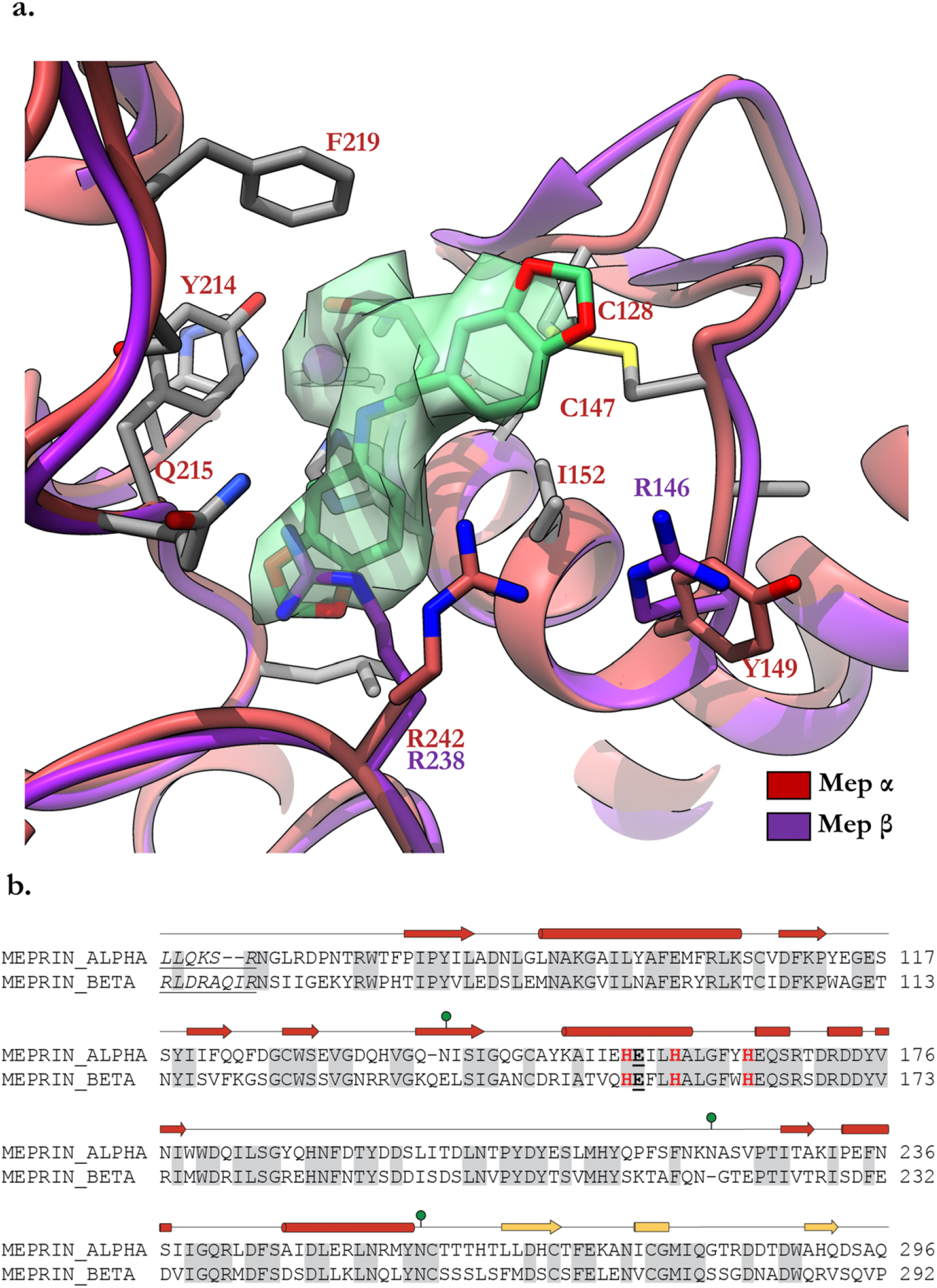
Structural comparison of meprin α and meprin β within the active site. **a.** Superposition of meprin α (red) and meprin β (purple) with select conserved side chains (grey) and divergent side chains (coloured) visible. **b.** Sequence alignment of meprin α and meprin β with secondary structure elements shown above (β-strands as arrows or α-helices as tubes). The catalytic glutamic acid is bold font and underlined. The active site tirade of histidines is shown in red bold font. Meprin α residue Y149 (meprin β R146) is non-conserved therefore presenting either as tyrosine or arginine in meprin α and meprin β respectively. This single residue appears to drive charge repulsion of R238 in meprin β (R242 in meprin α) causing a different rotamer position to be adopted relative to meprin α. This alternative rotamer conformation sterically occludes compound 10d (green; density shown as isosurface) suggesting how the drug remains selective despite major contacts of the drug and meprin α being conserved across both homologs. Conversely, in meprin α R242 is not repelled by Y149 and therefore adopts a conformation that does not interfere with drug binding.

## Bibliography

1 W. Scher. The role of extracellular proteases in cell proliferation and differentiation, 1987. ISSN 00236837.

2 Brigitte Bauvois. Transmembrane proteases in cell growth and invasion: New contributors to angiogenesis?, 2004. ISSN 09509232.

3 Marta Artal-Sanz and Nektarios Tavernarakis. Proteolytic mechanisms in necrotic cell death and neurodegeneration, 2005. ISSN 00145793.

4 Ting Jun Fan, Li Hui Han, Ri Shan Cong, and Jin Liang. Caspase family proteases and apoptosis. Acta Biochimica et Biophysica Sinica, 37(11):719–727, 2005. ISSN 16729145. doi: 10.1111/j.1745-7270.2005.00108.x.

5 Roopali Roy, Bo Zhang, and Marsha A. Moses. Making the cut: Protease-mediated regulation of angiogenesis, 2006. ISSN 00144827.

6 Katarina Wolf and Peter Friedl. Extracellular matrix determinants of proteolytic and non-proteolytic cell migration, 2011. ISSN 09628924.

7 Lijuan He and Denis Wirtz. Switching from protease-independent to protease-dependent cancercell invasion, 2014. ISSN 15420086.

8 Boris Turk. Targeting proteases: Successes, failures and future prospects, 2006. ISSN 14741776.

9 Ignacio Marín. Origin and diversification of meprin proteases. PLoS ONE, 10(8), 2015. ISSN 19326203. doi: 10.1371/journal.pone.0135924.

10 Erwin E. Sterchi, Walter Stöcker, and Judith S. Bond. Meprins, membrane-bound and secreted astacin metalloproteinases, 2008. ISSN 00982997.

11 Judith S. Bond and Robert J. Beynon. The astacin family of metalloendopeptidases, 1995. ISSN 1469896X.

12 R. J. Beynon, J. D. Shannon, and J. S. Bond. Purification and characterization of a metallo-endoproteinase from mouse kidney. The Biochemical journal, 199(3):591–598, 1981. ISSN 02646021. doi: 10.1042/bj1990591.

13 E. E. Sterchi, J. R. Green, and M. J. Lentze. Non-pancreatic hydrolysis of N-benzoyl-L-tyrosyl-p-aminobenzoic acid (PABA-peptide) in the human small intestine. Clinical Science, 62(5):557–560, 1982. ISSN 01435221. doi: 10.1042/cs0620557.

14 Judith S. Bond, Gail L. Matters, Sanjita Banerjee, and Renee E. Dusheck. Meprin metalloprotease expression and regulation in kidney, intestine, urinary tract infections and cancer, 2005. ISSN 00145793.

15 Christoph Becker-Pauly, Markus Höwel, Tatjana Walker, Annica Vlad, Karin Aufenvenne, Vinzenz Oji, Daniel Lottaz, Erwin E. Sterchi, Mekdes Debela, Viktor Magdolen, Heiko Traupe, and Walter Stöcker. The *α* and *β* subunits of the metalloprotease meprin are expressed in separate layers of human epidermis, revealing different functions in keratinocyte proliferation and differentiation. Journal of Investigative Dermatology, 127(5):1115–1125, 2007. ISSN 0022202X. doi: 10.1038/sj.jid.5700675.

16 Valentina Biasin, Leigh M. Marsh, Bakytbek Egemnazarov, Jochen Wilhelm, Bahil Ghanim, Walter Klepetko, Malgorzata Wygrecka, Horst Olschewski, Robert Eferl, Andrea Olschewski, and Grazyna Kwapiszewska. Meprin*β*, a novel mediator of vascular remodelling underlying pulmonary hypertension. Journal of Pathology, 233(1):7–17, 2014. ISSN 10969896. doi: 10.1002/path.4303.

17 Franka Scharfenberg, Fred Armbrust, Liana Marengo, Claus Pietrzik, and Christoph Becker-Pauly. Regulation of the alternative β-secretase meprin β by ADAM-mediated shedding, 2019. ISSN 14209071.

18 Tamara Jefferson, Ulrich Auf Dem Keller, Caroline Bellac, Verena V. Metz, Claudia Broder, Jana Hedrich, Anke Ohler, Wladislaw Maier, Viktor Magdolen, Erwin Sterchi, Judith S. Bond, Arumugam Jayakumar, Heiko Traupe, Athena Chalaris, Stefan Rose-John, Claus U. Pietrzik, Rolf Postina, Christopher M. Overall, and Christoph Becker-Pauly. The substrate degradome of meprin metalloproteases reveals an unexpected proteolytic link between meprin *β* and ADAM10. Cellular and Molecular Life Sciences, 70(2):309–333, 2013. ISSN 1420682X. doi: 10.1007/s00018-012-1106-2.

19 Daniel Kronenberg, Bernd C. Bruns, Catherine Moali, Sandrine Vadon-Le Goff, Erwin E. Sterchi, Heiko Traupe, Markus Böhm, David J.S. Hulmes, Walter Stöcker, and Christoph Becker-Pauly. Processing of procollagen III by meprins: New players in extracellular matrix assembly. Journal of Investigative Dermatology, 130(12):2727–2735, 2010. ISSN 15231747. doi: 10.1038/jid.2010.202.

20 Claudia Broder and Christoph Becker-Pauly. The metalloproteases meprin α and meprin β: Unique enzymes in inflammation, neurodegeneration, cancer and fibrosis, 2013. ISSN 02646021.

21 Johannes Prox, Philipp Arnold, and Christoph Becker-Pauly. Meprin *α* and meprin *β*: Procollagen proteinases in health and disease, 2015. ISSN 15691802.

22 Philipp Arnold, Inga Boll, Michelle Rothaug, Neele Schumacher, Frederike Schmidt, Rielana Wichert, Janna Schneppenheim, Juliane Lokau, Ute Pickhinke, Tomas Koudelka, Andreas Tholey, Björn Rabe, Jürgen Scheller, Ralph Lucius, Christoph Garbers, Stefan Rose-John, and Christoph Becker-Pauly. Meprin metalloproteases generate biologically active soluble interleukin-6 receptor to induce trans-signaling. Scientific Reports, 7, 2017. ISSN 20452322. doi: 10.1038/srep44053.

23 Tillmann Bedau, Florian Peters, Johannes Prox, Philipp Arnold, Frederike Schmidt, Malin Finkernagel, Sandra Köllmann, Rielana Wichert, Anna Otte, Anke Ohler, Marit Stirnberg, Ralph Lucius, Tomas Koudelka, Andreas Tholey, Valentina Biasin, Claus U. Pietrzik, Grazyna Kwapiszewska, and Christoph Becker-Pauly. Ectodomain shedding of CD99 within highly conserved regions is mediated by the metalloprotease meprin *β* and promotes transendothelial cell migration. FASEB Journal, 31(3):1226–1237, 2017. ISSN 15306860. doi: 10.1096/fj.201601113R.

24 V. Biasin, M. Wygrecka, L. M. Marsh, C. Becker-Pauly, L. Brcic, B. Ghanim, W. Klepetko, A. Olschewski, and G. Kwapiszewska. Meprin *β* contributes to collagen deposition in lung fibrosis. Scientific Reports, 7, 2017. ISSN 20452322. doi: 10.1038/srep39969.

25 Christian Herzog, Randy S. Haun, and Gur P. Kaushal. Role of meprin metalloproteinases in cytokine processing and inflammation, 2019. ISSN 10960023.

26 Florian Peters and Christoph Becker-Pauly. Role of meprin metalloproteases in metastasis and tumor microenvironment, 2019. ISSN 15737233.

27 Claudia Broder, Philipp Arnold, Sandrine Vadon Le Goff, Moritz A. Konerding, Kerstin Bahr, Stefan Müller, Christopher M. Overall, Judith S. Bond, Tomas Koudelka, Andreas Tholey, David J.S. Hulmes, Catherine Moali, and Christoph Becker-Pauly. Metalloproteases meprin *α* and meprin *β* are C-and N-procollagen proteinases important for collagen assembly and tensile strength. Proceedings of the National Academy of Sciences of the United States of America, 110(35):14219–14224, 2013. ISSN 10916490. doi: 10.1073/pnas.1305464110.

28 Christian Herzog, Gur P. Kaushal, and Randy S. Haun. Generation of biologically active interleukin-1*β* by meprin B. Cytokine, 31(5):394–403, 2005. ISSN 10434666. doi: 10.1016/j.cyto.2005.06.012.

29 Christian Herzog, Randy S. Haun, Varsha Kaushal, Philip R. Mayeux, Sudhir V. Shah, and Gur P. Kaushal. Meprin A and meprin *α* generate biologically functional IL-1 *β* from pro-IL-1*β*. Biochemical and Biophysical Research Communications, 379(4):904–908, 2009. ISSN 0006291X. doi: 10.1016/j.bbrc.2008.12.161.

30 Timothy R. Keiffer and Judith S. Bond. Meprin metalloproteases inactivate interleukin 6. JournalofBiologicalChemistry, 289(11):7580–7588, 2014. ISSN 1083351X. doi: 10.1074/jbc.M113.546309.

31 Sanjita Banerjee and Judith S. Bond. Prointerleukin-18 is activated by meprin *β* in vitro and in vivo in intestinal inflammation. Journal of Biological Chemistry, 283(46):31371–31377, 2008. ISSN 00219258. doi: 10.1074/jbc.M802814200.

32 Christian Herzog, Randy S. Haun, Sudhir V. Shah, and Gur P. Kaushal. Proteolytic processing and inactivation of CCL2/MCP-1 by meprins. Biochemistry and Biophysics Reports, 8: 146–150, 2016. ISSN 24055808. doi: 10.1016/j.bbrep.2016.08.019.

33 Christoph Becker-Pauly, Olivier Barré, Oliver Schilling, Ulrich Auf Dem Keller, Anke Ohler, Claudia Broder, André Schütte, Reinhild Kappelhoff, Walter Stöcker, and Christopher M. Overall. Proteomic analyses reveal an acidic prime side specificity for the astacin metalloprotease family reflected by physiological substrates. Molecular and Cellular Proteomics, 10(9), 2011. ISSN 15359476. doi: 10.1074/mcp.M111.009233.

34 Daniel Lottaz, Christoph A. Maurer, Dagmar Hahn, Markus W. Büchler, and Erwin E. Sterchi. Nonpolarized secretion of human meprin *α* in colorectal cancer generates an increased proteolytic potential in the stroma. Cancer Research, 59(5):1127–1133, 1999. ISSN 00085472.

35 Daniel Lottaz, Christoph A. Maurer, Agnès Noël, Silvia Blacher, Maya Huguenin, Alexandra Nievergelt, Verena Niggli, Alexander Kern, Stefan Müller, Frank Seibold, Helmut Friess, Christoph Becker-Pauly, Walter Stöcker, and Erwin E. Sterchi. Enhanced activity of meprin-α, a pro-migratory and pro-angiogenic protease, in colorectal cancer. PLoS ONE, 6(11), 2011. ISSN 19326203. doi: 10.1371/journal.pone.0026450.

36 André Schütte, Jana Hedrich, Walter Stöcker, and Christoph Becker-Pauly. Let it flow: Morpholino knockdown in zebrafish embryos reveals a pro-angiogenic effect of the metalloprotease meprin *α*2. PLoS ONE, 5(1), 2010. ISSN 19326203. doi: 10.1371/journal.pone.0008835.

37 Petra Minder, Elke Bayha, Christoph Becker-Pauly, and Erwin E. Sterchi. Meprin*α* transactivates the epidermal growth factor receptor (EGFR) via ligand shedding, thereby enhancing colorectal cancer cell proliferation and migration. Journal of Biological Chemistry, 287(42): 35201–35211, 2012. ISSN 00219258. doi: 10.1074/jbc.M112.368910.

38 Emilie Vazeille, Marie Agnès Bringer, Aurélie Gardarin, Christophe Chambon, Christoph Becker-Pauly, Sylvia L.F. Pender, Christine Jakob, Stefan Müller, Daniel Lottaz, and Arlette Darfeuille-Michaud. Role of meprins to protect ileal Mucosa of Crohn’s disease patients from colonization by adherent-invasive e. coli. PLoS ONE, 6(6), 2011. ISSN 19326203. doi: 10.1371/journal.pone.0021199.

39 Rielana Wichert, Anna Ermund, Stefanie Schmidt, Matthias Schweinlin, Miroslaw Ksiazek, Philipp Arnold, Katharina Knittler, Frederike Wilkens, Barbara Potempa, Björn Rabe, Marit Stirnberg, Ralph Lucius, Jörg W. Bartsch, Susanna Nikolaus, Maren Falk-Paulsen, Philip Rosenstiel, Marco Metzger, Stefan Rose-John, Jan Potempa, Gunnar C. Hansson, Peter J. Dempsey, and Christoph Becker-Pauly. Mucus Detachment by Host Metalloprotease Meprin *β* Requires Shedding of Its Inactive Pro-form, which Is Abrogated by the Pathogenic Protease RgpB. Cell Reports, 21(8):2090–2103, 2017. ISSN 22111247. doi: 10.1016/j.celrep.2017.10.087.

40 Daniel Ramsbeck, Antje Hamann, Dagmar Schlenzig, Stephan Schilling, and Mirko Buchholz. First insight into structure-activity relationships of selective meprin *β* inhibitors. Bioorganic and Medicinal Chemistry Letters, 27(11):2428–2431, 2017. ISSN 14643405. doi: 10.1016/j.bmcl.2017.04.012.

41 Daniel Ramsbeck, Antje Hamann, Georg Richter, Dagmar Schlenzig, Stefanie Geissler, Vera Nykiel, Holger Cynis, Stephan Schilling, and Mirko Buchholz. Structure-Guided Design, Synthesis, and Characterization of Next-Generation Meprin *β* Inhibitors. Journal of Medicinal Chemistry, 61(10):4578–4592, 2018. ISSN 15204804. doi: 10.1021/acs.jmedchem.8b00330.

42 Kathrin Tan, Christian Jäger, Dagmar Schlenzig, Stephan Schilling, Mirko Buchholz, and Daniel Ramsbeck. Tertiary-Amine-Based Inhibitors of the Astacin Protease Meprin *α*. Chem Med Chem, 13(16):1619–1624, 2018. ISSN 18607187. doi: 10.1002/cmdc.201800300.

43 Christoph Becker, Markus N. Kruse, Kristina A. Slotty, Danny Köhler, J. Robin Harris, Sandra Rösmann, Erwin E. Sterchi, and Walter Stöcker. Differences in the activation mechanism between the *α* and *β* subunits of human meprin. Biological Chemistry, 384(5): 825–831, 2003. ISSN 14316730. doi: 10.1515/BC.2003.092.

44 Petra Marchand, Jie Tang, and Judith S. Bond. Membrane association and oligomeric organization of the *α* and *β* subunits of mouse meprin A. Journal of Biological Chemistry, 269(21):15388–15393,1994. ISSN 00219258. doi: 10.1016/s0021-9258(17)36618-8.

45 Jeremy A. Hengst and Judith S. Bond. Transport of meprin subunits through the secretory pathway: Role of the transmembrane and cytoplasmic domains and oligomerization. Journal of Biological Chemistry, 279(33):34856–34864, 2004. ISSN 00219258. doi: 10.1074/jbc.M405774200.

46 Eric Dumermuth, Joyce A. Eldering, Jürgen Grünberg, Weiping Jiang, and Erwin E. Sterchi. Cloning of the PABA peptide hydrolase alpha subunit (PPH*α*) from human small intestine and its expression in COS-1 cells. FEBS Letters, 335(3):367–375, 1993. ISSN 00145793. doi: 10.1016/0014-5793(93)80421-P.

47 Christian Herzog, Randy S. Haun, Andreas Ludwig, Sudhir V. Shah, and Gur P. Kaushal. ADAM10 is the major sheddase responsible for the release of membrane-Associated meprinA. Journal of Biological Chemistry, 289(19):13308–13322,2014. ISSN 1083351X. doi: 10.1074/jbc.M114.559088.

48 Florian Peters, Franka Scharfenberg, Cynthia Colmorgen, Fred Armbrust, Rielana Wichert, Philipp Arnold, Barbara Potempa, Jan Potempa, Claus U. Pietrzik, Robert Häsler, Philip Rosenstiel, and Christoph Becker-Pauly. Tethering soluble meprin *α* in an enzyme complex to the cell surface affects IBD-associated genes. FASEB Journal, 33(6):7490–7504, 2019. ISSN 15306860. doi: 10.1096/fj.201802391R.

49 Sylvie Chevallier, Jinhi Ahn, Guy Boileau, and Philippe Crine. Identification of the cysteine residues implicated in the formation of *α*2 and *α*/*β* dimers of rat meprin. Biochemical Journal, 317(3):731–738, 1996. ISSN 02646021. doi: 10.1042/bj3170731.

50 Greg P Bertenshaw, Mona T. Norcum, and Judith S. Bond. Structure of homo-and heterooligomeric meprin metalloproteases: Dimers, tetramers, and high molecular mass multimers. Journal of Biological Chemistry, 278(4):2522–2532, 2003. ISSN 00219258. doi: 10.1074/jbc.M208808200.

51 Faoud T. Ishmael, Mona T. Norcum, Stephen J. Benkovic, and Judith S. Bond. Multimeric Structure of the Secreted Meprin A Metalloproteinase and Characterization of the Functional Protomer. Journal of Biological Chemistry, 276(25):23207–23211, 2001. ISSN 00219258. doi: 10.1074/jbc.M102654200.

52 J G Tate, S Bamford, H C Jubb, Z Sondka, D M Beare, N Bindal, H Boutselakis, C G Cole, C Creatore, E Dawson, P Fish, B Harsha, C Hathaway, S C Jupe, C Y Kok, K Noble, L Ponting, C C Ramshaw, C E Rye, H E Speedy, R Stefancsik, S L Thompson, S Wang, S Ward, P J Campbell, and S A Forbes. COSMIC: the Catalogue Of Somatic Mutations In Cancer. Nucleic Acids Res, 47(D1):D941–D947, 2019. ISSN 1362-4962 (Electronic) 0305-1048 (Linking). doi: 10.1093/nar/gky1015.

53 Joan L. Arolas, Claudia Broder, Tamara Jefferson, Tibisay Guevara, Erwin E. Sterchi, Wolfram Bode, Walter Stöcker, Christoph Becker-Pauly, and F. Xavier Gomis-Rüth. Structural basis for the sheddase function of human meprin *β* metalloproteinase at the plasma membrane. Proceedings of the National Academy of Sciences of the United States of America, 109(40):16131–16136, 2012. ISSN 00278424. doi: 10.1073/pnas.1211076109.

54 Miriam Linnert, Claudia Fritz, Christian Jäger, Dagmar Schlenzig, Daniel Ramsbeck, Martin Kleinschmidt, Michael Wermann, Hans Ulrich Demuth, Christoph Parthier, and Stephan Schilling. Structure and dynamics of meprin *β* in complex with a hydroxamate-based inhibitor. International Journal of Molecular Sciences, 22(11), 2021. ISSN 14220067. doi: 10.3390/ijms22115651.

55 Julien Robert-Paganin, Xiao Ping Xu, Mark F. Swift, Daniel Auguin, James P. Robblee, Hailong Lu, Patricia M. Fagnant, Elena B. Krementsova, Kathleen M. Trybus, Anne Houdusse, Niels Volkmann, and Dorit Hanein. The actomyosin interface contains an evolutionary conserved core and an ancillary interface involved in specificity. Nature Communications, 12 (1),2021. ISSN 20411723. doi: 10.1038/s41467-021-22093-4.

56 Matthieu P.M.H. Benoit, Ana B. Asenjo, and Hernando Sosa. Cryo-EM reveals the structural basis of microtubule depolymerization by kinesin-13s. Nature Communications, 9(1), 2018. ISSN 20411723. doi: 10.1038/s41467-018-04044-8.

57 C. Herzog, R. Seth, S. V. Shah, and G. P. Kaushal. Role of meprin a in renal tubular epithelial cell injury. Kidney International, 71(10):1009–1018, 2007. ISSN 00852538. doi: 10.1038/sj.ki.5002189.

58 Haim Ashkenazy, Elana Erez, Eric Martz, Tal Pupko, and Nir Ben-Tal. ConSurf 2010: Calculating evolutionary conservation in sequence and structure of proteins and nucleic acids. Nucleic Acids Research, 2010. ISSN 03051048. doi: 10.1093/nar/gkq399.

59 Haim Ashkenazy, Shiran Abadi, Eric Martz, Ofer Chay, Itay Mayrose, Tal Pupko, and Nir Ben-Tal. ConSurf 2016: an improved methodology to estimate and visualize evolutionary conservation in macromolecules. Nucleic acids research, 2016. ISSN 13624962. doi: 10.1093/nar/gkw408.

60 Markus N. Kruse, Christoph Becker, Daniel Lottaz, Danny Köhler, Irene Yiallouros, Hans Willi Krell, Erwin E. Sterchi, and Walter Stöcker. Human meprin *α* and *β* homooligomers: Cleavage of basement membrane proteins and sensitivity to metalloprotease inhibitors. Biochemical Journal, 378(2):383–389, 2004. ISSN 02646021. doi: 10.1042/BJ20031163.

61 S. S. Craig, J. F. Reckelhoff, and J. S. Bond. Distribution of meprin in kidneys from mice with high-and low-meprin activity. American Journal of Physiology - Cell Physiology, 253 (4), 1987. ISSN 00029513. doi: 10.1152/ajpcell.1987.253.4.c535.

62 Howard Trachtman, Elsa Valderrama, Janet M. Dietrich, and Judith S. Bond. The role of meprin A in the pathogenesis of acute renal failure. Biochemical and Biophysical Research Communications, 208(2):498–505, 1995. ISSN 0006291X. doi: 10.1006/bbrc.1995.1366.

63 Gur P. Kaushal, Randy S. Haun, Christian Herzog, and Sudhir V. Shah. Meprin A metalloproteinase and its role in acute kidney injury, 2013. ISSN 03636127.

64 David S. Goodsell and Arthur J. Olson. Structural symmetry and protein function, 2000. ISSN 10568700.

65 Mayssam H. Ali and Barbara Imperiali. Protein oligomerization: How and why, 2005. ISSN 09680896.

66 Birgitta Tomkinson. Tripeptidyl-peptidase II: Update on an oldie that still counts, 2019. ISSN 61831638.

67 Birgitta Tomkinson. Association and dissociation of the tripeptidyl-peptidase II complex as a way of regulating the enzyme activity. Archives of Biochemistry and Biophysics, 376(2): 275–280, 2000. ISSN 00039861. doi: 10.1006/abbi.2000.1713.

68 Birgitta Tomkinson, Bairbre Ní Laoi, and Kimberly Wellington. The insert within the catalytic domain of tripeptidyl-peptidase II is important for the formation of the active complex. European Journal of Biochemistry, 269(5):1438–1443, 2002. ISSN 00142956. doi: 10.1046/j.1432-1033.2002.02783.x.

69 Laёtitia Gorisse, Christine Pietrement, Vincent Vuiblet, Christian E.H. Schmelzer, Martin Köhler, Laurent Duca, Laurent Debelle, Paul Fornès, Stéphane Jaisson, and Philippe Gillery. Protein carbamylation is a hallmark of aging. Proceedings of the National Academy of Sciences of the United States of America, 113(5):1191–1196, 2016. ISSN 10916490. doi: 10.1073/pnas.1517096113.

70 David N. Mastronarde. Automated electron microscope tomography using robust prediction of specimen movements. Journal of Structural Biology, 152(1):36–51, 2005. ISSN 10478477. doi: 10.1016/j.jsb.2005.07.007.

71 James R. Kremer, David N. Mastronarde, and J. Richard McIntosh. Computer visualization of three-dimensional image data using IMOD. Journal of Structural Biology, 116(1):71–76, 1996. ISSN 10478477. doi: 10.1006/jsbi.1996.0013.

72 Shawn Q Zheng, Eugene Palovcak, Jean-Paul Armache, Kliment A Verba, Yifan Cheng, and David A Agard. MotionCor2: anisotropic correction of beam-induced motion for improved cryo-electron microscopy. Nature Methods, 14(4):331–332, apr 2017. ISSN 1548-7091. doi: 10.1038/nmeth.4193.

73 David N. Mastronarde and Susannah R. Held. Automated tilt series alignment and tomographic reconstruction in IMOD. Journal of Structural Biology, 197(2):102–113, 2017. ISSN 10958657. doi: 10.1016/j.jsb.2016.07.011.

74 David N. Mastronarde. Dual-axis tomography: An approach with alignment methods that preserve resolution. In Journal of Structural Biology, volume 120, pages 343–352, 1997. doi: 10.1006/jsbi.1997.3919.

75 Beata Turonová, Florian K.M. Schur, William Wan, and John A.G. Briggs. Efficient 3D-CTF correction for cryo-electron tomography using NovaCTF improves subtomogram averaging resolution to 3.4 Å. Journal of Structural Biology, 199(3):187–195, 2017. ISSN 10958657. doi: 10.1016/j.jsb.2017.07.007.

76 Daniel Castaño-Díez, Mikhail Kudryashev, Marcel Arheit, and Henning Stahlberg. Dynamo: A flexible, user-friendly development tool for subtomogram averaging of cryo-EM data in high-performance computing environments. Journal of Structural Biology, 2012. ISSN 10478477. doi: 10.1016/j.jsb.2011.12.017.

77 Alexis Rohou and Nikolaus Grigorieff. CTFFIND4: Fast and accurate defocus estimation from electron micrographs. Journal of Structural Biology, 192(2):216–221, nov 2015. ISSN 1047-8477. doi: 10.1016/J.JSB.2015.08.008.

78 Dimitry Tegunov and Patrick Cramer. Real-time cryo-electron microscopy data preprocessing with Warp. Nature Methods, 2019. ISSN 15487105. doi: 10.1038/s41592-019-0580-y.

79 Thorsten Wagner, Felipe Merino, Markus Stabrin, Toshio Moriya, Claudia Antoni, Amir Apelbaum, Philine Hagel, Oleg Sitsel, Tobias Raisch, Daniel Prumbaum, Dennis Quentin, Daniel Roderer, Sebastian Tacke, Birte Siebolds, Evelyn Schubert, Tanvir R. Shaikh, Pascal Lill, Christos Gatsogiannis, and Stefan Raunser. SPHIRE-crYOLO: A fast and accurate fully automated particle picker for cryo-EM. Comms Biol, 2:218(2019):1–13, 2019. ISSN 2399-3642. doi: 10.1101/356584.

80 Thorsten Wagner, Luca Lusnig, Sabrina Pospich, Markus Stabrin, Fabian Schonfeld, and Stefan Raunser. Two particle-picking procedures for filamentous proteins: SPHIRE-crYOLO filament mode and SPHIRE-STRIPER. Acta Crystallographica Section D: Structural Biology, 76:613–620, 2020. ISSN 20597983. doi: 10.1107/S2059798320007342.

81 Sjors H.W. Scheres. RELION: Implementation of a Bayesian approach to cryo-EM structure determination. Journal of Structural Biology, 180(3):519–530, 2012. ISSN 10478477. doi: 10.1016/j.jsb.2012.09.006.

82 Takanori Nakane, Erik Lindahl, Jasenko Zivanov, Wim JH Hagen, Sjors HW Scheres, Dari Kimanius, and Björn O Forsberg. New tools for automated high-resolution cryo-EM structure determination in RELION-3. eLife, 7, nov 2018. ISSN 2050-084X. doi: 10.7554/elife.42166.

83 Jasenko Zivanov, Takanori Nakane, and Sjors H.W. Scheres. Estimation of high-order aberrations and anisotropic magnification from cryo-EM data sets in RELION-3.1. IUCr J, 2020. ISSN 20522525. doi: 10.1107/S2052252520000081.

84 Ali Punjani, John L Rubinstein, David J Fleet, and Marcus A Brubaker. cryoSPARC: algorithms for rapid unsupervised cryo-EM structure determination. Nature Methods, (August 2016), 2017. doi: 10.1038/nmeth.4169.

85 Ali Punjani, Haowei Zhang, and David J. Fleet. Non-uniform refinement: adaptive regularization improves single-particle cryo-EM reconstruction. Nature Methods, 17(12):1214– 1221, 2020. ISSN 15487105. doi: 10.1038/s41592-020-00990-8.

86 Ali Punjani and David J Fleet. 3D Flexible Refinement: Structure and Motion of Flexible Proteins from Cryo-EM. bioRxiv, page 2021.04.22.440893, jan 2021. doi: 10.1101/2021.04.22.440893.

87 Serban L Ilca, Abhay Kotecha, Xiaoyu Sun, Minna M Poranen, David I Stuart, and Juha T Huiskonen. Localized reconstruction of subunits from electron cryomicroscopy images of macromolecular complexes. Nature Communications, 6:1–8, 2015. doi: 10.1038/ncomms9843.

88 R Sanchez-Garcia, J Gomez-Blanco, A Cuervo, J M Carazo, C O S Sorzano, and J Vargas. DeepEMhancer: a deep learning solution for cryo-EM volume post-processing. bioRxiv, page 2020.06.12.148296, jan 2020. doi: 10.1101/2020.06.12.148296.

89 D. Asarnow, E. Palovcak, and Y. Cheng. UCSF pyem, 2019.

90 Andrew Waterhouse, Martino Bertoni, Stefan Bienert, Gabriel Studer, Gerardo Tauriello, Rafal Gumienny, Florian T. Heer, Tjaart A.P. De Beer, Christine Rempfer, Lorenza Bordoli, Rosalba Lepore, and Torsten Schwede. SWISS-MODEL: Homology modelling of protein structures and complexes. Nucleic Acids Research, 46(W1):W296–W303, 2018. ISSN 13624962. doi: 10.1093/nar/gky427.

91 Eric F. Pettersen, Thomas D. Goddard, Conrad C. Huang, Gregory S. Couch, Daniel M. Greenblatt, Elaine C. Meng, and Thomas E. Ferrin. UCSF Chimera - A visualization system for exploratory research and analysis. Journal of Computational Chemistry, 25(13):1605– 1612, 2004. ISSN 01928651. doi: 10.1002/jcc.20084.

92 Tristan Ian Croll. ISOLDE: A physically realistic environment for model building into low-resolution electron-density maps. Acta Crystallographica Section D: Structural Biology, 74 (6):519–530, 2018. ISSN 20597983. doi: 10.1107/S2059798318002425.

93 D Goddard Thomas, C Huang Conrad, C Meng Elaine, F Pettersen Eric, S Couch Gregory, H Morris John, and E Ferrin Thomas. UCSF ChimeraX: Meeting modern challenges in visualization and analysis. Protein Science, 27(1):14–25, 2017. ISSN 0961-8368. doi: 10.1002/pro.3235.

94 P Emsley and K Cowtan. Coot: model-building tools for molecular graphics. Acta Crystal-logrDBiolCrystallogr, 60(Pt 12 Pt 1):2126–2132, 2004. ISSN 0907-4449 (Print) 0907-4449 (Linking). doi: 10.1107/S0907444904019158.

95 P. Emsley, B. Lohkamp, W. G. Scott, and K. Cowtan. Features and development of Coot. Acta Crystallographica Section D: Biological Crystallography, 66(4):486–501, 2010. ISSN 09074449. doi: 10.1107/S0907444910007493.

96 Ulrich Eckhard, Hagen Körschgen, Nele von Wiegen, Walter Stöcker, and F. Xavier Gomis-Rüth. The crystal structure of a 250-kDa heterotetrameric particle explains inhibition of sheddase meprin *β* by endogenous fetuin-B. Proceedings of the National Academy of Sciences of the United States of America, 118(14), 2021. ISSN 10916490. doi: 10.1073/pnas.2023839118.

97 John Jumper, Richard Evans, Alexander Pritzel, Tim Green, Michael Figurnov, Olaf Ronneberger, Kathryn Tunyasuvunakool, Russ Bates, Augustin Žídek, Anna Potapenko, Alex Bridgland, Clemens Meyer, Simon A. A. Kohl, Andrew J. Ballard, Andrew Cowie, Bernardino Romera-Paredes, Stanislav Nikolov, Rishub Jain, Jonas Adler, Trevor Back, Stig Petersen, David Reiman, Ellen Clancy, Michal Zielinski, Martin Steinegger, Michalina Pacholska, Tamas Berghammer, Sebastian Bodenstein, David Silver, Oriol Vinyals, Andrew W. Senior, Koray Kavukcuoglu, Pushmeet Kohli, and Demis Hassabis. Highly accurate protein structure prediction with AlphaFold. Nature, jul 2021. ISSN 0028-0836. doi: 10.1038/s41586-021-03819-2.

98 Mihaly Varadi, Stephen Anyango, Mandar Deshpande, Sreenath Nair, Cindy Natassia, Galabina Yordanova, David Yuan, Oana Stroe, Gemma Wood, Agata Laydon, Augustin Žídek, Tim Green, Kathryn Tunyasuvunakool, Stig Petersen, John Jumper, Ellen Clancy, Richard Green, Ankur Vora, Mira Lutfi, Michael Figurnov, Andrew Cowie, Nicole Hobbs, Pushmeet Kohli, Gerard Kleywegt, Ewan Birney, Demis Hassabis, and Sameer Velankar. AlphaFold Protein Structure Database: massively expanding the structural coverage of protein-sequence space with high-accuracy models. Nucleic Acids Research, 50(D1): D439–D444, jan 2022. ISSN 0305-1048. doi: 10.1093/nar/gkab1061.

99 Pavel V. Afonine, Billy K. Poon, Randy J. Read, Oleg V. Sobolev, Thomas C. Terwilliger, Alexandre Urzhumtsev, Paul D. Adams, and IUCr. Real-space refinement in *PHENIX* for cryo-EM and crystallography. Acta Crystallographica Section D Structural Biology, 74 (6):531–544, jun 2018. ISSN 2059-7983. doi: 10.1107/S2059798318006551.

100 P D Adams, P V Afonine, G Bunkoczi, V B Chen, I W Davis, N Echols, J J Headd, L W Hung, G J Kapral, R W Grosse-Kunstleve, A J McCoy, N W Moriarty, R Oeffner, R J Read, D C Richardson, J S Richardson, T C Terwilliger, and P H Zwart. PHENIX: a comprehensive Python-based system for macromolecular structure solution. Acta Crystallogr D Biol Crystallogr, 66(Pt 2):213–221, 2010. ISSN 1399-0047 (Electronic) 0907-4449 (Linking). doi: 10.1107/S0907444909052925.

101. 101 Min Chen, Mohamed R. Daha, and Cees G M Kallenberg. The complement system in systemic autoimmune disease. Journal of Autoimmunity, 34(3):J276–J286, 2010. ISSN 08968411. doi: 10.1016/j.jaut.2009.11.014.

