## Supplementary Table 1 for "Helical ultrastructure of the oncogenic metalloprotease meprin α in complex with a small molecule hydroxamate inhibitor"

**Supp Table 1 Cryo-EM data collection, refinement and validation statistics**

|  | **Pro-Meprin α**  (EMD-26419)  (PDB 7UAB)  **Dataset-180314** | **Pro-Meprin α**  (EMD-26420)  (PDB 7UAC)  **Dataset-190220** | **Meprin α**  **(helix)**  (EMD-26421)  **Dataset-190220** | **Meprin α**  (EMD-26422)  (PDB 7UAE)  **Dataset-190220** | **Meprin α / Compound 10d**  (EMD-26423)  (PDB 7UAF)  **Dataset-190218** | **Meprin α /**  **Fetuin-B**  **(helix)**  (EMD-26424)  **Dataset-220125** | **Meprin α /**  **Fetuin-B**  (EMD-26426)  (PDB 7UAI)  **Dataset-220125** |
| --- | --- | --- | --- | --- | --- | --- | --- |
| **Data collection and processing** |  |  |  |  |  |  |  |
| Magnification | 130,000× | 130,000× | 130,000× | 130,000× | 130,000× | 130,000× | 130,000× |
| Voltage (kV) | 300 | 300 | 300 | 300 | 300 | 300 | 300 |
| Electron exposure  (e^–^/Å^2^) | 22.25 | 44.5 | 44.5 | 44.5 | 44.5 | 44.5 | 44.5 |
| Defocus range (μm) | −0.5 to −2.2 | −0.5 to −1.5 | −0.5 to −1.5 | −0.5 to −1.5 | −0.5 to −1.2 | −0.5 to −2.0 | −0.5 to −2.0 |
| Pixel size (Å)(binned) | 1.06 | 1.06 | 1.06 | 1.06 | 1.06 | 1.06 (1.4133) | 1.06 (1.4133) |
| Symmetry imposed | C1 | C1 | C1 | C1 | C1 | C1 | C1 |
| Initial particle images (no.) | 580,400 | 905,660 | 147,823 | 905,660 | 925,458 | 132,975 | 519,760 |
| Final particle images (no.) | 116,080 | 338,429 | 111,312 | 233,978 | 235,162 | 103,952 | 115,951 |
| Map resolution (Å) | 3.7 / 4.1 | 2.7 / 3.3 | 3.4 / 3.9 | 2.6 / 3.0 | 2.4 / 2.8 | 3.7 / 4.2 | 2.8 / 3.7 |
| 0.143/0.5 FSC threshold |  |  |  |  |  |  |  |
| Map resolution range (Å) | 3.6 - 7.0 | 2.8 – 4.2 | 3.3 - 18.0 | 2.6 – 3.9 | 2.4 – 4.6 | 3.5 – 22.2 | 2.9 – 8.1 |
| 0.5 FSC threshold |  |  |  |  |  |  |  |
| 3DFSC sphericity | 0.98 | 0.96 | 0.87 | 0.97 | 0.98 | 0.86 | 0.98 |
| 0.5 FSC threshold |  |  |  |  |  |  |  |
| Reconstruction type | Subparticle localised | Subparticle + 3D masked refinement | Consensus | Subparticle + 3D masked refinement | Subparticle localised | Consensus | Subparticle localised |
| **Refinement** |  |  |  |  |  |  |  |
| Initial model  (PDB code) | 4GWN / AF-Q16819-v2 | 7UAB |  | 7UAB | 4GWN |  | AF-Q9UGM5-v2, 7UAB, 7AUW |
| Model resolution (Å)  0.5 FSC threshold | 3.8 | 3.0 |  | 2.8 | 2.7 |  | 3.4 |
| Map sharpening *B* factor* (Å^2^) | −85 | −103 |  | −94 | −57 |  | −54 |
| Model composition |  |  |  |  |  |  |  |
| Non-hydrogen atoms | 16,877 | 4,540 |  | 4,513 | 17,864 |  | 21,938 |
| Protein residues | 2,042 | 546 |  | 535 | 2,131 |  | 2,690 |
| *B* factors (Å^2^) |  |  |  |  |  |  |  |
| Protein | 80.38 | 85.46 |  | 71.06 | 39.92 |  | 64.97 |
| Ligands | 99.52 | 100.47 |  | 94.18 | 61.79 |  | 91/12 |
| R.M.S. deviations |  |  |  |  |  |  |  |
| Bond lengths (Å) | 0.012 | 0.012 |  | 0.011 | 0.012 |  | 0.013 |
| Bond angles (°) | 1.722 | 1.781 |  | 1.742 | 1.801 |  | 1.777 |
| Validation |  |  |  |  |  |  |  |
| MolProbity score | 1.09 | 0.88 |  | 0.82 | 1.01 |  | 1.16 |
| Clashscore | 1.07 | 0.57 |  | 0.11 | 1.19 |  | 1.32 |
| Poor rotamers (%) | 0.44 | 0.00 |  | 0.00 | 0.16 |  | 0.90 |
| Ramachandran plot |  |  |  |  |  |  |  |
| Favored (%) | 95.96 | 96.84 |  | 96.06 | 96.93 |  | 95.82 |
| Allowed (%) | 3.99 | 3.16 |  | 3.94 | 3.07 |  | 4.24 |
| Disallowed (%) | 0.05 | 0.00 |  | 0.00 | 0.00 |  | 0.04 |

*As determined by the Rosenthal and Henderson method, however for model building amplitude correction and map sharpening was sometimes performed by deepEMhancer which does not generate a *B*-factor.
